# Multi-omic analysis of lung tumors defines pathways activated in neuroendocrine transformation

**DOI:** 10.1101/2020.12.02.408476

**Authors:** Alvaro Quintanal-Villalonga, Hirozaku Taniguchi, Yingqian A. Zhan, Maysun M. Hasan, Fanli Meng, Fathema Uddin, Mark Donoghue, Helen H. Won, Shweta S. Chavan, Joseph M. Chan, Metamia Ciampricotti, Andrew Chow, Michael Offin, Jason C. Chang, Jordana Ray-Kirton, Jacklynn Egger, Umesh K. Bhanot, Joachim Silber, Christine A. Iacobuzio-Donahue, Michael H. Roehrl, Travis J. Hollmann, Helena A. Yu, Natasha Rekhtman, John T. Poirier, Brian Houck-Loomis, Richard P. Koche, Charles M. Rudin, Triparna Sen

## Abstract

Lineage plasticity, a capacity to reprogram cell phenotypic identity under evolutionary pressure, is implicated in treatment resistance and metastasis in multiple cancers. In lung adenocarcinomas (LUADs) amenable to treatment with targeted inhibitors, transformation to an aggressive neuroendocrine (NE) carcinoma resembling small cell lung cancer (SCLC) is a recognized mechanism of acquired resistance. Defining molecular mechanisms of NE transformation in lung cancer has been limited by a paucity of well annotated pre- and post-transformation clinical samples. We hypothesized that mixed histology LUAD/SCLC tumors may capture cancer cells proximal to, and on either side of, histologic transformation. We performed detailed genomic, epigenomic, transcriptomic and proteomic characterization of combined LUAD/SCLC tumors as well as pre- and post-transformation clinical samples. Our data support that NE transformation is primarily driven by transcriptional reprogramming rather than mutational events. We identify genomic contexts in which NE transformation is favored, including frequent loss of the 3p chromosome arm in pre-transformation LUADs. Consistent shifts in gene expression programs in NE transformation include induction of several stem/progenitor cell regulatory pathways, including upregulation of PRC2 and WNT signaling, and suppression of Notch pathway activity. We observe induction of PI3K/AKT and an immunosuppressive phenotype in NE transformation. Taken together our findings define a novel landscape of potential drivers and therapeutic vulnerabilities of NE transformation in lung cancer.

## INTRODUCTION

Lineage plasticity describes the capacity of cells to transition from one committed identity to that of a distinct developmental lineage. This phenotypic flexibility can promote survival of cancer cells under unfavorable conditions, such as hypoxia or selective pressure from oncogenic driver-targeted therapy^1,2,3^.

The histological transformation of lung adenocarcinoma (LUAD) to an aggressive neuroendocrine (NE) derivative resembling small cell lung cancer (SCLC) is a signature example of lineage plasticity in cancer. Transformed SCLC (T-SCLC) is associated with a notably poor prognosis, similar or worse than that of *de novo* SCLC^3^. Initially described in the prostate setting as a mechanism of resistance to androgen suppression^4–6^, this phenomenon was then identified in LUADs harboring *EGFR* mutations^7^, and subsequently found to occur more broadly in lung cancers^8^. The increased practice of tumor re-biopsy upon disease progression has improved the ability to identify histologic transformation, which in *EGFR*-mutant LUAD may comprise up to 14% of cases of acquired resistance to osimertinib^9,10^.

Identification of the molecular mechanisms promoting lineage plasticity in clinical samples is key to identifying patients at high risk of transformation and may define strategies to prevent or treat this phenomenon. Little is known about the molecular alterations occurring during NE transformation in human tumors, including in lung cancer. Transcriptomic analyses of prostate cancer undergoing NE histologic transformation have been performed, but only on relapsed and post-transformation samples^11,12^. A paucity of well-annotated paired pre- and post-transformation clinical samples has been a major hurdle in defining mechanisms of lineage plasticity in lung cancer. Previous genomic studies in small numbers of cases, have suggested that concomitant inactivation of *TP53* and *RB1* is necessary but not sufficient, and have reported other recurrent genomic alterations^3,9,13^.

On rare occasions, pathologic examination of resected cancers reveals more than one histology in single tumors. We hypothesized that such cases might represent lineage plasticity captured in temporal and spatial proximity to the occurrence of a histologic shift. Detailed molecular characterization of such cases could provide novel insight into key drivers of histologic transformation. Here we report the first comprehensive characterization of NE transformation, including genomic, transcriptomic, epigenomic and proteomic analyses, in a cohort of mixed histology LUAD/SCLC samples. In addition to our primary analysis of mixed histology tumors with discrete areas of LUAD and SCLC, we include analyses in matched pre- and post-transformation cases, with reference to control “pure” LUAD and SCLC. Our strategy provides novel insights into molecular drivers and potential therapeutic vulnerabilities of NE transformation in lung cancer.

## RESULTS

### Genomic landscape defines novel predictors of NE transformation

For in-depth characterization of NE transformation, we analyzed clinical specimens consisting of combined LUAD/SCLC histology exhibiting clear spatial separation (n=11); pre-transformation LUADs (n=5) and post-transformation SCLCs (n=3), including one matched case; never-transformed LUADs (n=15); and *de novo* SCLCs (n=18) (**Figure 1A and Supplementary Tables S1-S4**). Microdissection was performed for independent genomic, epigenomic, transcriptomic, and immunohistochemical analyses (**Figure 1B, 1C and Supplementary Figure S1**).

**Figure 1.**
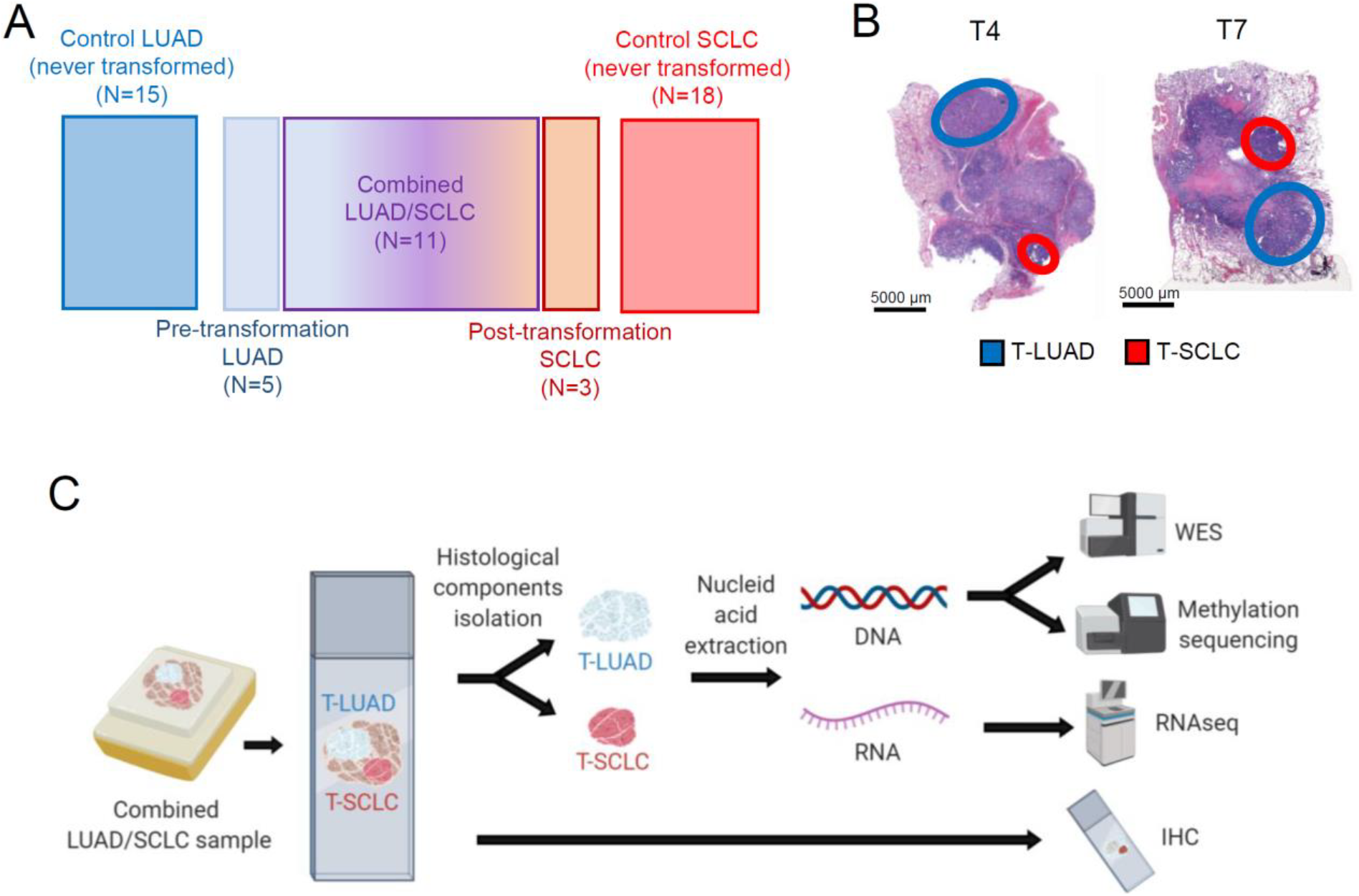
Multilayer molecular characterization of SCLC transformation. Related to Supplementary Figure S1. (A) Schematic composition of the cohort under study. (B) Illustrative H&E images of two of our combined histology samples, showing spatial separation of both independently isolated histologic components. (C) Schema of processing of combined histology samples for molecular analyses.

Our selection of combined histology samples for this analysis was predicated on the assumption that the LUAD and SCLC components were clonally related. Alternatively, it was possible that these represented “collision tumors” derived from two independent oncogenic events. Whole exome sequencing (WES) of all LUAD and SCLC samples from combined histology specimens and the matched pre- and post-transformation pair (T12) revealed multiple shared mutations in all cases, confirming that matched LUAD and SCLC components were clonal (**Figure 2A**). We therefore refer to these hereafter as T-LUAD and T-SCLC with the T referring to histologic transformation, without presumption of directionality. Higher tumor purity in the T-SCLC component was observed, consistent with the low stromal content of SCLC relative to LUAD^14,15^ (**Supplementary Figure S2A**). We did not observe consistent differences in tumor ploidy, tumor mutation burden, or predicted neoantigen burden between T-LUAD and T-SCLC components (**Supplementary Figures S2B-D**).

**Figure 2.**
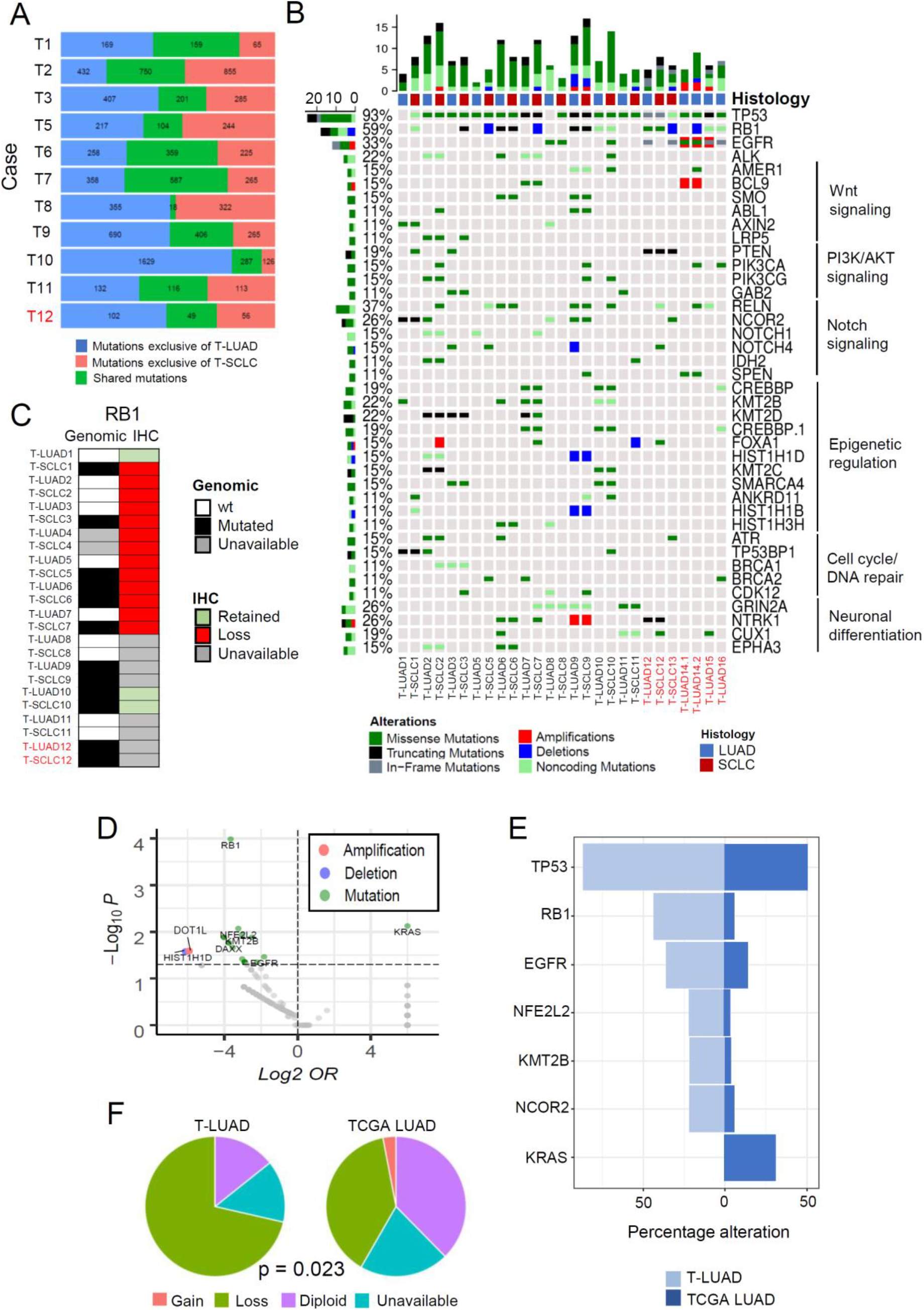
Genomic characterization of SCLC transformation. Related to Supplementary Figures S2-3. (A) Bar blot showing number of mutations occurring specifically in the T-LUAD and T-SCLC components, and of mutations shared between these. (B) Oncoprint showing the most prevalent mutations and CNAs in the transformation samples, grouped by recurrent pathways. (C) Heatmap showing complementary genomic and immunohistochemical characterization of RB1 alterations. (D) Volcano plot showing enrichment of mutations/CNAs in T-LUAD versus TCGA LUAD cohort. (E) Bar plot showing prevalence (%) of mutations/CNA enriched in T-LUAD versus TCGA LUAD with over 25% prevalence in our cohort. (F) Pie charts showing the abundance of 3p chromosome arm lost in our T-LUAD cases versus TCGA LUAD. p-value for enrichment in 3p loss was calculated using the Fisher’s exact test for count data. Samples IDs in black and red indicate that they come from a combined histology specimen or a pre-/post-transformation specimen, respectively.

We next sought to define mutational processes that might contribute to lineage plasticity and histologic transformation through mutational signature analysis. Smoking signature was dominant in 7 out of 11 cases but did not differ consistently between T-LUAD and T-SCLC (**Supplementary Figure S2E**). APOBEC signature, previously proposed to be a predictor of SCLC-transformation in triple *EGFR/TP53/RB1* mutant tumors^9^, was prominent only 5 out of 11 of the T-LUAD samples (**Supplementary Figure S2E**).

Analyses of the most prevalent mutations and copy number alterations (CNAs), including variants of both known and unknown significance, revealed almost universal *TP53* loss in both T-LUAD and T-SCLC (93%, **Figure 2B**), with only two T-LUADs (T-LUAD1 and T-LUAD8) showing wild type *TP53*. *RB1* mutations/deletions were less frequently detected (59% of samples), identified in 7 out of 14 T-LUADs, and in 8 out of 11 T-SCLCs (**Figures 2B, C**). However, IHC in samples for which tissue was available showed that Rb protein expression was lost in all but one T-LUAD (T-LUAD1) and in all T-SCLC samples (**Figure 2C**). These results show that loss of *RB1* function might be independent of genomic alterations, highlighting the importance of complementary genomic and IHC profiling for confirmation of *RB1* activity. Oncogenic *EGFR* mutations were present in 33% of T-LUAD samples (**Figure 2B**), further illustrating that NE transformation may occur outside the *EGFR* mutant setting^8^. Within matched pairs, we observed common mutations of both known and unknown significance highlighting genetic relatedness. There were no recurrent mutational events seen in more than two cases in this dataset, suggesting that while a preexisting genetic context may facilitate plasticity, NE transformation itself may not be mutationally driven.

To better define the context that may permit lineage plasticity, we focused on the most commonly altered genes in this sample set, present in both the T-LUAD and T-SCLC components (**Figure 2B**). Notably, these include factors involved in WNT signaling (*BCL9, SMO, AXIN2*, etc.); PI3K/AKT signaling (*PTEN, PIK3CA, PIK3CG*, etc.); Notch signaling (*NOTCH1/4, SPEN, RELN*, etc.); epigenetic regulation (*KMT2B/C/D, CREBBP, SMARCA4* and *FOXA1*); cell cycle/DNA repair (*ATR, BRCA1/2* and *TP53BP)*; and neural differentiation (*NTRK1, CUX1, GRIN2A*). The presence of these pathway alterations in T-LUAD samples implies that they may occur early in the NE transformation process and may prime LUAD for lineage transition.

Next, we compared the frequency of mutations/copy number alteration (CNAs) identified in the T-LUADs in our cohort to those of the TCGA LUADs (**Figures 2D,E and Supplementary Figure S3**). We focused on the differentially mutated genes showing alterations in ≥ 20% of T-LUAD samples, to filter for those more likely to have a role in transformation promotion. As expected, we found enrichment of *TP53* (p=0.013) and *RB1* (p<0.001) alterations in T-LUAD^3,9^. Consistent with previous reports of NE transformation in *EGFR*-mutant LUAD, we found enrichment in *EGFR* alterations (p=0.035) in the T-LUAD cohort^9,13^. We noted decreased frequency of *KRAS* mutations in our T-LUAD (p=0.008); this may suggest that *KRAS*-mutant LUADs are less likely to undergo NE transformation, or, alternatively, may be attributable to the historical lack of potent targeted inhibitors of KRAS.

Novel observations in this analysis included mutations on *NFE2L2* (p=0.009), a transcription factor involved in response to oxidative stress^16^; *KMT2B* (p=0.012), an epigenetic regulator; and *NCOR2* (p=0.045), a transcriptional corepressor involved in Notch signaling^17^. These genes were altered in almost 25% of T-LUADs, but only rarely (<5%) in the TGCA LUADs (**Figure 2E and supplementary Figure S3A**). Validation in a larger cohort would be required to support a potential role for *NFE2L2, KMT2B* and *NCOR2* alterations as predictors of susceptibility to SCLC transformation. Interestingly, we observed recurrent loss of the 3p chromosome arm in ~75% of our pre-transformation LUAD cases, a significantly higher rate than observed in TCGA LUADs (p=0.023, **Figure 2F and Supplementary Figure S3B**). Chromosome 3p arm loss may comprise a novel predictive biomarker for SCLC transformation.

### Clonal evolution of SCLC transformation

The paired nature of the combined histology tumors provided an opportunity to explore serial events in branched evolution of the distinct histologic lineages. WES data was of sufficient quality to allow reconstruction of the clonal history for 5 of the cases under study (**Figure 3**). For each of these, we identified exclusive or enriched mutations in the T-SCLC components, but no common mutations across cases were observed. Additionally, we observed numerous copy number alterations (**Figure 3, left**), potentially caused by genomic instability derived from *TP53* mutations, which were identified as an early event in most samples. Many of these were shared within each matched pair (**Figure 3, left**), suggesting that they occurred before the histologic divergence. Interestingly, these analyses also suggested that the whole genome doubling events observed in each of these cases occurred after the clonal split leading to NE lineage commitment (**Figure 3, right**).

**Figure 3.**
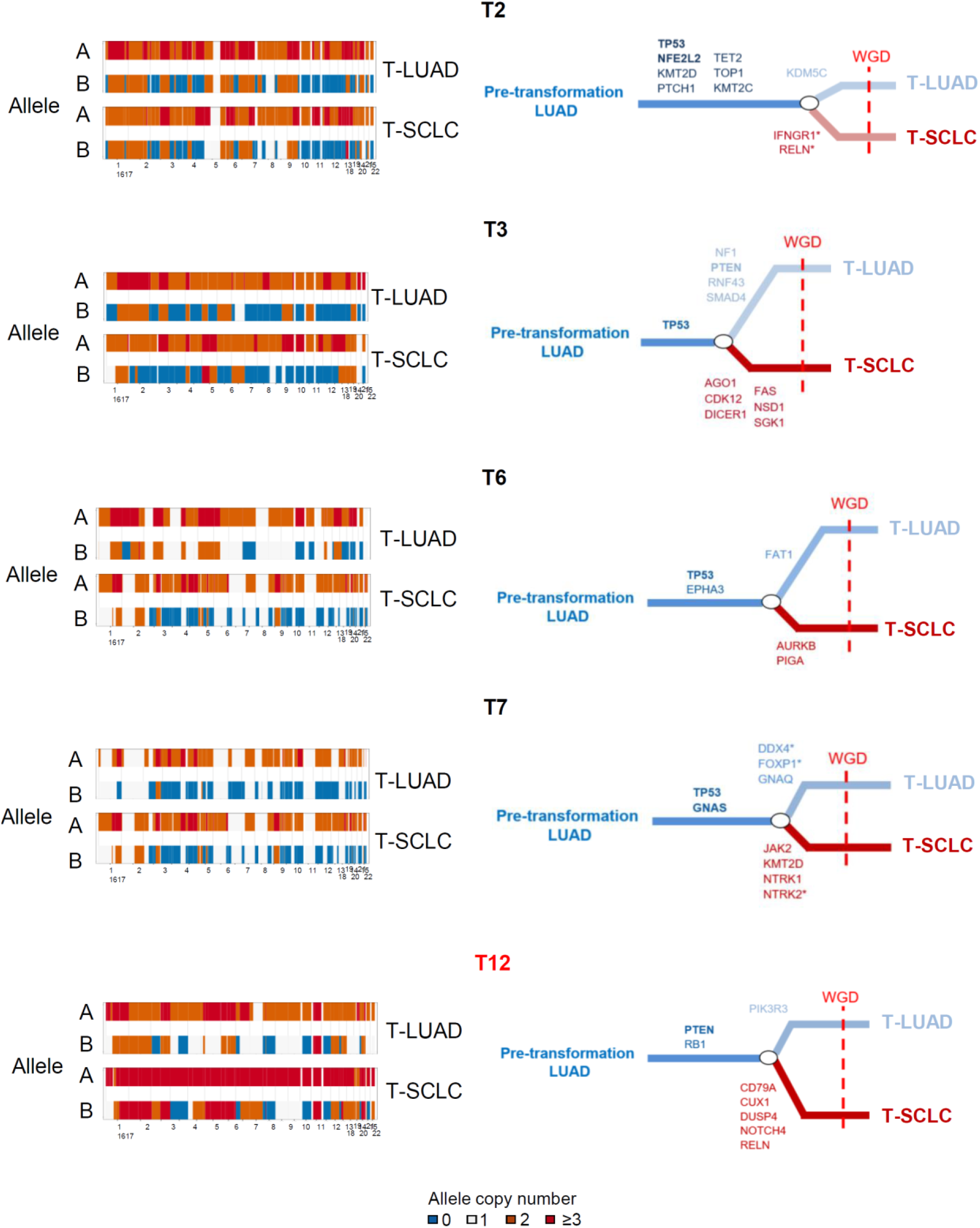
Clonal mutation evolution of SCLC transformation. Chromosomal gain/losses in both alleles for matched LUAD and SCLC components for each case (left) and reconstruction of clonal evolution (right) in 4 combined histology and 1 pair of pre- and post-transformation cases. Whole genome doubling (WGD) event is indicated by a red dashed line. Genes in bold letter are indicative of the occurrence of a hotspot mutation. Genes with an asterisk (“*”) indicate the presence of that particular mutation in the other histological component at subclonal level. Samples IDs in black and red indicate that they come from a combined histology specimen or a pre-/post-transformation specimen, respectively.

### T-SCLC spans all SCLC subtypes

Work from our group and others has highlighted the inter-tumoral heterogeneity of SCLCs^15^. *De novo* SCLCs can be divided into discrete molecularly defined subtypes based on dominant expression of one of four transcriptional regulators: *ASCL1* (SCLC-A), *NEUROD1* (SCLC-N), *POU2F3* (SCLC-P), and *YAP1* (SCLC-Y). However, little is known about the molecular subtyping of T-SCLC tumors, or whether these tumors consistently align with one of these four defined subtypes.

To study if T-SCLCs were enriched in any subtype, we analyzed relative expression of these four transcriptional regulators in the T-LUADs and T-SCLC samples at both mRNA and protein (IHC) levels (**Figure 4A and Supplementary Table S5)**. Expression of three transcription factors (*ASCL1*, *NEUROD1* and *POU2F3*) was consistently low in the T-LUADs. However, expression of *YAP1* was higher in all but one (T4) T-LUADs than in their matched T-SCLCs (**Figure 4A**). *YAP1* expression was higher in never-transformed LUADs that T-LUAD (**Supplementary Figure S4A**), consistent with the oncogenic role of this Hippo pathway effector in LUAD^18,19^ and with its incompatibility with NE features in lung cancer^20^. We observed good concordance between IHC and RNA data. Where discrepant, we assigned the subtype based on relative RNA expression, following current consensus^15^.

**Figure 4.**
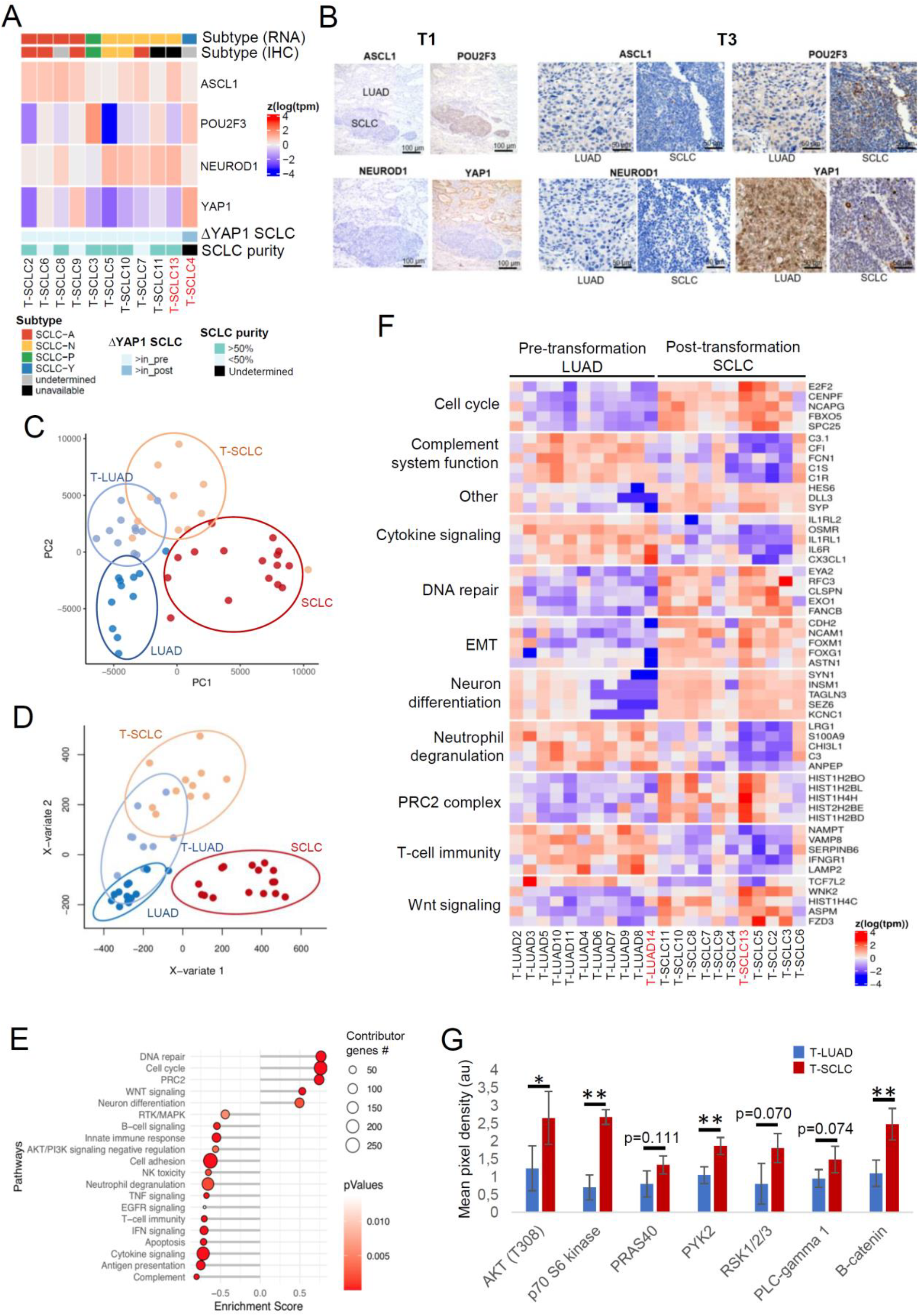
Transcriptomic, epigenomic and proteomic characterization of SCLC transformation. Related to Supplementary Figures S4-6. (A) Heatmap showing mRNA expression of the SCLC subtype-determining TFs, tumor purity, highest TF expressed by IHC in the T-SCLC component and YAP1 mRNA expression in the T-SCLC component relative to their matched T-LUAD component, in the transformation samples. (B) IHC images for subtype-determining TFs in the SCLC-P T-SCLC cases (ch1 and ch3). (C) PCA analysis on the transcriptomes of our pre- and post-transformation samples, and of our control LUAD and *de novo* SCLC samples. (D) PLSDA analyses on the methylome of our T-LUAD and T-SCLC samples, and of our control LUAD and SCLC samples. (E) Pathway enrichment analyses on the DEGs of the T-LUAD versus T-SCLC comparison. (F) Heatmap highlighting DEGs of interest, grouped by recurrent pathways. (G) Bar plot showing differential phosphorylation of genes involved in the AKT/Wnt signaling pathways, and differential expression of β-catenin, as determined by an antibody array on pre- and post-transformation clinical and PDX samples. Samples IDs in black and red indicate that they come from a combined histology specimen or a pre-/post-transformation specimen, respectively. p-values legend: * p<0.05, **p<0.01.

Notably, we were able to detect all four SCLC subtypes among the T-SCLC samples, suggesting that lineage plasticity in LUAD can give rise to any of the four SCLC subtypes. Interestingly, two of the samples (T1 and T3) were categorized as SCLC-P, with high *POU2F3* levels exclusive to the T-SCLC component, and no expression of any of the other transcription factors by IHC (**Figures 4A,B and Supplementary Table S5**). Tuft cells, a rare population of lung cells, have been previously hypothesized to be the cell of origin for SCLC-P, based on a very similar POU2F3-dependent gene expression program exclusive in this cell type of the normal lung^21^. However, no POU2F3 protein expression was observed in the matched T-LUAD components of these samples (**Figures 4A,B and Supplementary Table S5**). Furthermore, mRNA levels of other tuft cell markers^21^ were also not elevated in these T-LUADs relative to the rest of pre-transformation or control LUADs (**Supplementary Figure S4B**). This suggests that a tuft cell-like gene expression program is induced in this T-SCLC subtype, independent of the cell of origin. Hence our results, for the first time, demonstrate that T-SCLCs conform to all major subtypes of *de novo* SCLCs, and suggest a tuft cell-independent origin of SCLC-P.

### Gene expression and methylation analyses identify pathways involved in NE transformation

We performed transcriptomic (RNAseq) and methylation (EPIC) analyses of T-LUADs, T-SCLCs, control LUADs and *de novo* SCLCs (**Figure 1C and Supplementary Tables S1-3**). Principal component analyses (PCA) of the RNAseq data showed dissimilar expression patterns for control LUAD and *de novo* SCLC, as expected (**Figure 4C**). T-LUADs clustered together in adjacency to control LUADs, and T-SCLCs in proximity to *de novo* SCLCs. T-LUAD and T-SCLC did appear to represent intermediate phenotypes, and demonstrated substantial overlap in expression profile (**Figure 4C**). This suggests that T-LUADs might be distinctly primed to transform, relative to other LUAD, and that T-SCLC retains some transcriptomic features of T-LUAD. PCA analysis of methylation profiling by EPIC revealed that T-SCLCs exhibit distinct methylation profiles to those of *de novo* SCLCs, and show proximity to the methylome of T- and control LUADs (**Figure 4D and Supplementary Figures S4C**). This implies that tumors undergoing NE transformation retain broad scale epigenomic features of the LUAD from which they derived.

To further analyze the transcriptional changes occurring during NE transformation, we performed differential gene expression and pathway enrichment analyses (GSEA) of T-LUAD and T-SCLC samples (**Figure 4E**). As expected, T-SCLC demonstrated increased expression of NE markers such as *SYP, SYN1* and *INSM1*; and genes associated to Notch signaling inhibition, such as *DLL3* and *HES6*^15^. Pathway enrichment analyses performed on differentially expressed genes (DEG) in T-LUAD vs. T-SCLC samples (**Figures 4E,F**) showed T-SCLC-specific upregulation of genes involved in (1) neural differentiation (including *SEZ6, TAGLN3* and *KCNC1)*; (2) cell cycle progression (including *E2F2, CENPF* and *FBXO5)*; (3) DNA repair (including *FANCB, EYA2* and *RFC3*); (4) chromatin remodeling (including *HDAC2*); and (5) PRC2 complex (including *HIST1H2BO, HIST1H2BL and HISH1H4H*) (**Figures 4E,F**). We further confirmed a consistent increase in the mRNA expression of *EZH2,* one of the main components of the PRC2 complex (**Supplementary Figure S5A**), previously strongly implicated in lineage plasticity and neuroendocrine transformation in prostate cancer^4^. GSEA analyses also showed a gene expression signature of induced WNT signaling in T-SCLC, with downregulation of the negative regulator of WNT signaling *TCF7L2* and overexpression of WNT pathway activators such as *WNK2, ASPM and FZD3* (**Figures 4E-F**). This was further confirmed at the protein level by protein arrays of T-LUAD and T-SCLC samples and patient-derived xenografts (PDXs) (**Figure 4G**). We observed increased expression of the major WNT signaling effector, β-catenin, and increased phosphorylation of PYK2, a protein involved in WNT signaling activation^22^ in T-SCLC (**Figure 4G and Supplementary Table S7**). Among other changes, NE transformation was also associated with global downregulation of receptor tyrosine kinase signaling, inhibition of apoptotic induction, suppression of anti-tumor immune activation, and induction of PI3K/AKT signaling (**Figures 4E,F,G)**.

### Integration of gene expression and DNA methylation data

Integrative analyses of transcriptomic and epigenomic data showed that a substantial number of differentially expressed genes were also differentially methylated in T-SCLC relative to T-LUAD, consistent with epigenomic reprogramming upon NE transformation in lung. We observed cell adhesion, neuron differentiation, cytokine signaling and neutrophil degranulation pathways to be among the top pathways differentially affected by methylation (**Figure 5A and Supplementary Figure S5C**).

**Figure 5.**
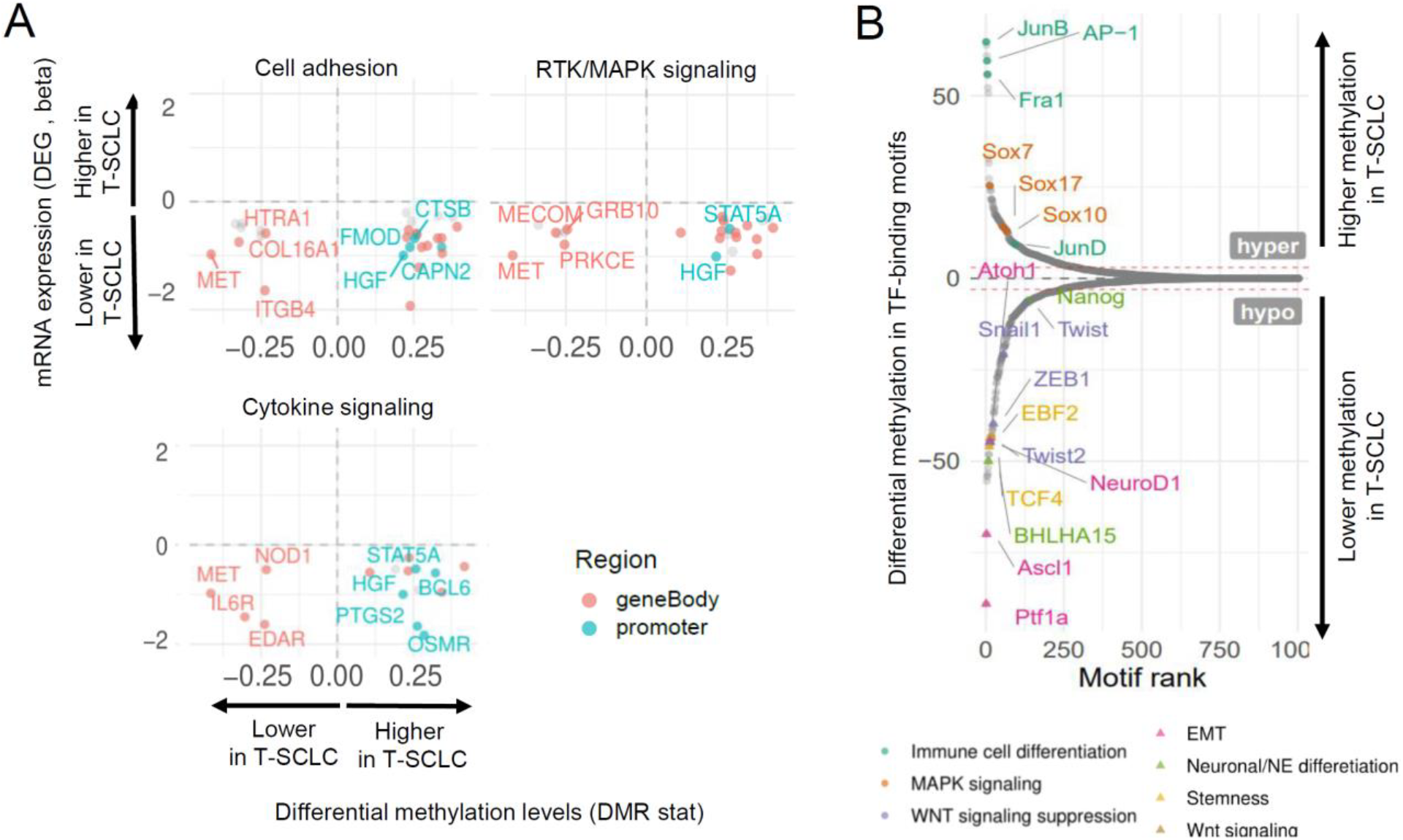
Integrative RNA and methylation analyses of SCLC transformation. Related to Supplementary Figure S7. (A) Scatter plots showing DEGs exhibiting differential methylation levels in T-LUAD versus control LUAD comparison, grouped by pathways of interest. Significantly differentially expressed (q value < 0.05 and beta >= log2(1.5)) and methylated (FDR < 0.5 and differential methylation level greater than 0.1) sites are highlighted. Those genes where increased gene body or promoter methylation is correlated to expression positively and negatively, respectively, are labeled. (B) Plot exhibiting differentially methylated transcription factor binding domains in T-SCLC versus T-LUAD. Interested TFs in this study are highlighted and labeled.

Methylation occurring in TF-binding motifs can inhibit TF binding and affect regulation of target gene expression^23^. Analysis of differential methylation of TF-binding motifs revealed hypomethylation of binding motifs for genes involved in (1) neuronal and NE differentiation (including *ASCL1* and *NEUROD1)*; (2) WNT signaling activators (*TCF4, EBF2*); (3) stemness (*NANOG, BHLHA15*); and (4) EMT (*SNAI1, TWIST1/2, ZEB1*, among others) in T-SCLC relative to T-LUAD (**Figure 5B**). We also found T-SCLC-specific hypermethylation of binding motifs for TFs involved in MAPK signaling (*JUNB/D, AP-1, FOSL1/2*); and WNT signaling suppression (*SOX7/10/17*) (**Figure 5B**). These data suggest that epigenomic reprogramming upon transformation leads to altered methylation of key TF-binding motifs, driving expression phenotypes observed during histological transition (**Figures 4E-F**).

Notably, three TFs, *FOXN4* (β=3.38, q-value=0.031), *ONECUT2* (β=3.10, q-value=0.014) and *POU3F2* (β=2.02, q-value=0.083), were among the top differentially expressed genes upregulated in T-SCLCs (**Supplementary Figure S6A**). ONECUT2 and POU3F2 have been previously implicated in acquisition and maintenance of the neuroendocrine phenotype in prostate cancer^24,25^. FOXN4 has been previously shown to interact with ASCL1 to modulate Notch signaling^26^. To assess the role of these TFs as drivers of NE transformation, we overexpressed *FOXN4, ONECUT2,* and *POU3F2* each independently in two *EGFR*-mutant LUAD cell lines (PC9 and HCC827, **Supplementary Figures S6B-C**). Ectopic overexpression of these factors did not induce upregulation of neuroendocrine markers at the protein level (ASCL1, NEUROD1, chromogranin A, synaptophysin; data not shown), but did downregulate EGFR expression in both lines (**Supplementary Figure S6B**). These results suggest that although these transcription factors may not individually be key effectors of NE transformation *per se*, they might be involved through downregulation of EGFR expression, a commonly observed phenotype in EGFR-mutant SCLC transformed samples^3,13^.

Taken together, these data highlight that while epigenetic reprogramming in NE transformation results in induction of transcriptional changes affecting several key signaling pathways, some epigenomic features are maintained during NE transformation, differentiating these tumors from *de novo* SCLC. Transformation to a neuroendocrine phenotype may be promoted by the PRC2 complex and other epigenetic modifiers, and appears to be characterized by activation of PI3K and WNT signaling pathways, acquisition of a mesenchymal phenotype, and suppression of anti-tumor immune response pathways.

### Transcriptomic and epigenomic analyses of T-LUADs reveal early molecular alterations in NE transformation

To identify transcriptional changes that may predispose to NE transformation, we next compared the transcriptomic and methylomic profiles of T-LUAD and control (never-transformed) LUADs (**Figures 6A-C**). In the T-LUAD samples, we observed relative downregulation of a variety of keratin genes (*KRT7, KRT8* and *KRT15*, among others, **Figure 6C**), consistent with a potential partial loss of LUAD phenotype^27^. As expected, we also observed multiple alterations in the RB pathway (**Figure 6A**). *RB1* mutations and Rb protein loss were found in 36% (4/11) and in 86% (6/7), respectively, of T-LUADs. We also observed differential expression of members upstream *RB1* (possibly in compensation for *RB1* functional deficiency^28,29^) including upregulation of *CDKN2A* associated with an increases in gene body methylation (**Figures 6A and Supplementary Figure S7A**); downregulation of *CCND1* (Cyclin D1) and upregulation of *CCNE1/2* (Cyclin E1/2). These results are consistent with prior observations that *RB1* loss of function precedes NE transformation^3,9^.

**Figure 6.**
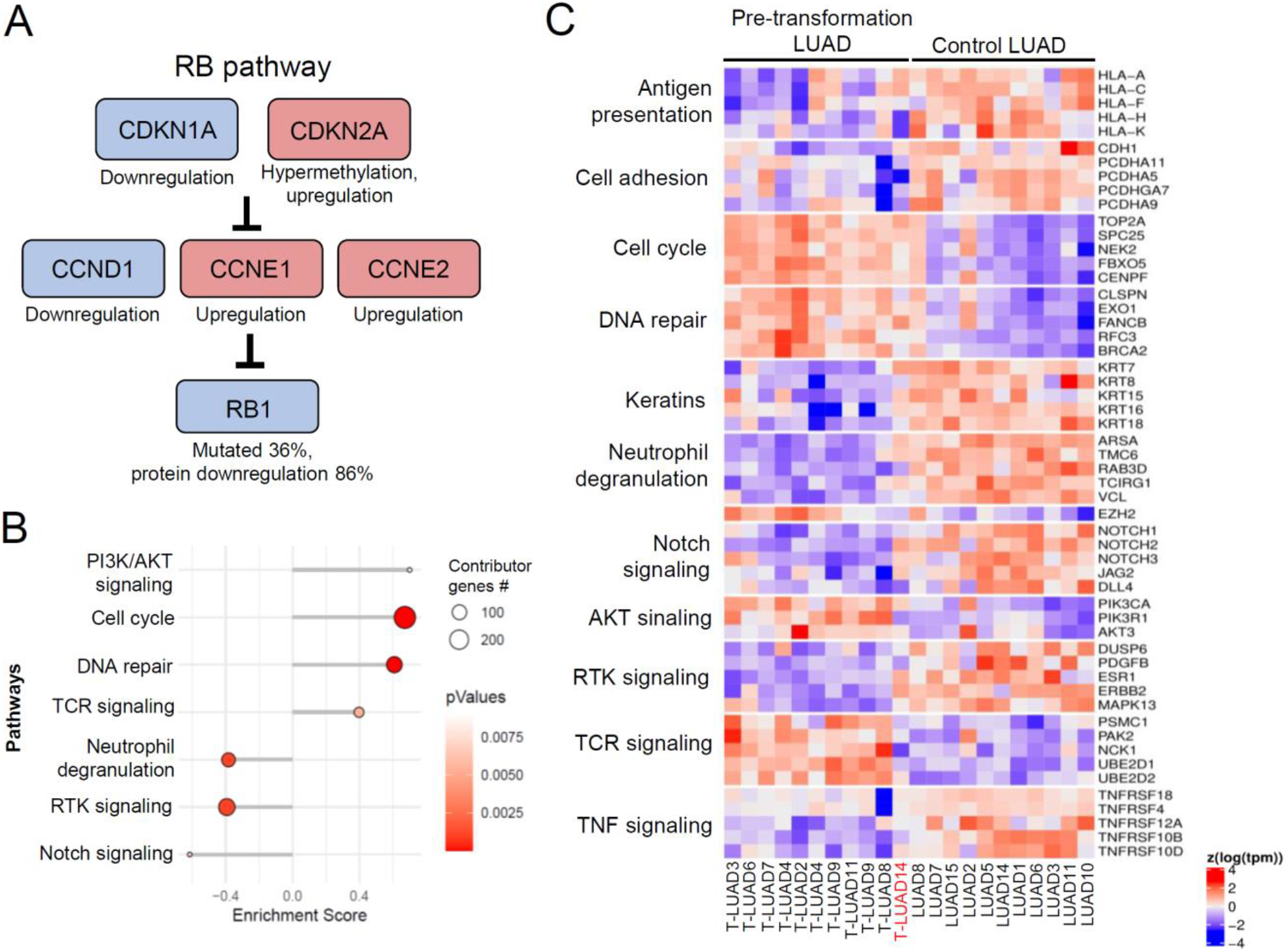
Integrative RNA and methylation analyses of T-LUAD versus control LUAD. Related to Supplementary Figure S7 (A) Alterations in the RB pathway identified in T-LUAD. (B) Pathway enrichment analyses on the DEGs of the T-LUAD versus control LUAD comparison. (C) Heatmap highlighting DEGs of interest, grouped by recurrent pathways, of the T-LUAD versus control LUAD comparison. Samples IDs in black and red indicate that they come from a combined histology specimen or a pre-/post-transformation specimen, respectively.

We also identified DEGs representative of some of the same pathways identified when comparing T-LUAD and T-SCLC samples, suggesting progressive differential regulation of these pathways in NE transformation (**Figures 6B-C**). These included up-regulation of genes enriched in cell cycle progression (*TOP2A, CENPF, FBXO5*), DNA repair pathways (*CLSPN, EXO1, FANCB*), and PI3K/AKT signaling (*PIK3CA, PIK3R1, AKT3*); as well as downregulation of RTK signaling (*DUSP6, ERBB2,* and *MAPK13*), cell adhesion (*CDH1* (E-cadherin)*, PCDHA11, PCDHA9*) and anti-tumor immune response (multiple genes involved in neutrophil degranulation, TNF signaling and antigen presentation). Consistent with the known role of Notch signaling in suppressing NE tumor growth^30^, these analyses revealed early downregulation of genes involved in Notch signaling, including Notch receptors *NOTCH1/2/3*, and ligands *JAG2* and *DLL4* (**Figures 6B-C**). Consistent with an overall retention of genome methylation patterns of LUAD, integrative analyses with transcriptomic and methylation data revealed that none of these pathways was likely being differentially regulated by gene-specific methylation (**Supplementary Figure S7B**).

These results suggest that an intermediate phenotype is captured in T-LUAD specimens, which is further accentuated upon NE transformation to T-SCLC. This phenotype is characterized by partial loss of LUAD features and of dependence on RTK signaling, and by the upregulation of gene programs promoting AKT signaling, cell cycle progression and DNA repair, as well as downregulation of genes related to immune response and Notch signaling.

### Molecular comparison of de novo and T-SCLCs reveals differential signaling and immune pathways regulation

Finally, we sought to explore molecular differences between transformed and *de novo* SCLCs. Comparison of the transcriptome of T-SCLCs to that of our control *de novo* SCLCs revealed lower expression of genes involved in neuron differentiation (*SALL3, DLX1,* and *NEURL1*); Notch signaling (*JAG2, DLL1/4,* and *NOTCH3*); PI3K/AKT pathway (*AKT1/2, BAD*, and *TSC2*); and epigenetic regulators (*HIST2H3D, SMARCA4* and *ARID1B*) (**Figures 7A-B**). We also observed higher expression of genes involved in stemness (such as *CD44, NAMPT* or the aldehyde dehydrogenase *ALDH1A2)*; IFN signaling (*TLR2/3/7/8, CLEC7A*), lymphocyte chemotaxis (*CXCL10/13/14, XCL* and , *CCL5*) and TCR signaling (*PAK2, UBE2D2,* and *NCK1*) in T-SCLCs relative to de novo SCLCs. Integrative transcriptome/methylome analyses (**Figure 7C and Supplementary Figure S7C**) indicated that the suppressed neuronal phenotype in T-SCLCs was associated with a high number of differentially methylated genes in that pathway, suggesting epigenetic reprogramming (**Figure 7C**). These results suggest that T-SCLC may be characterized by decreased neuronal features, an accentuated stem-like/plastic phenotype, and increased ability to induce an anti-tumor immune response relative to *de novo* SCLC. These data further support that inhibition of Notch signaling may be particularly key for SCLC transformation and persists after histological transition.

**Figure 7.**
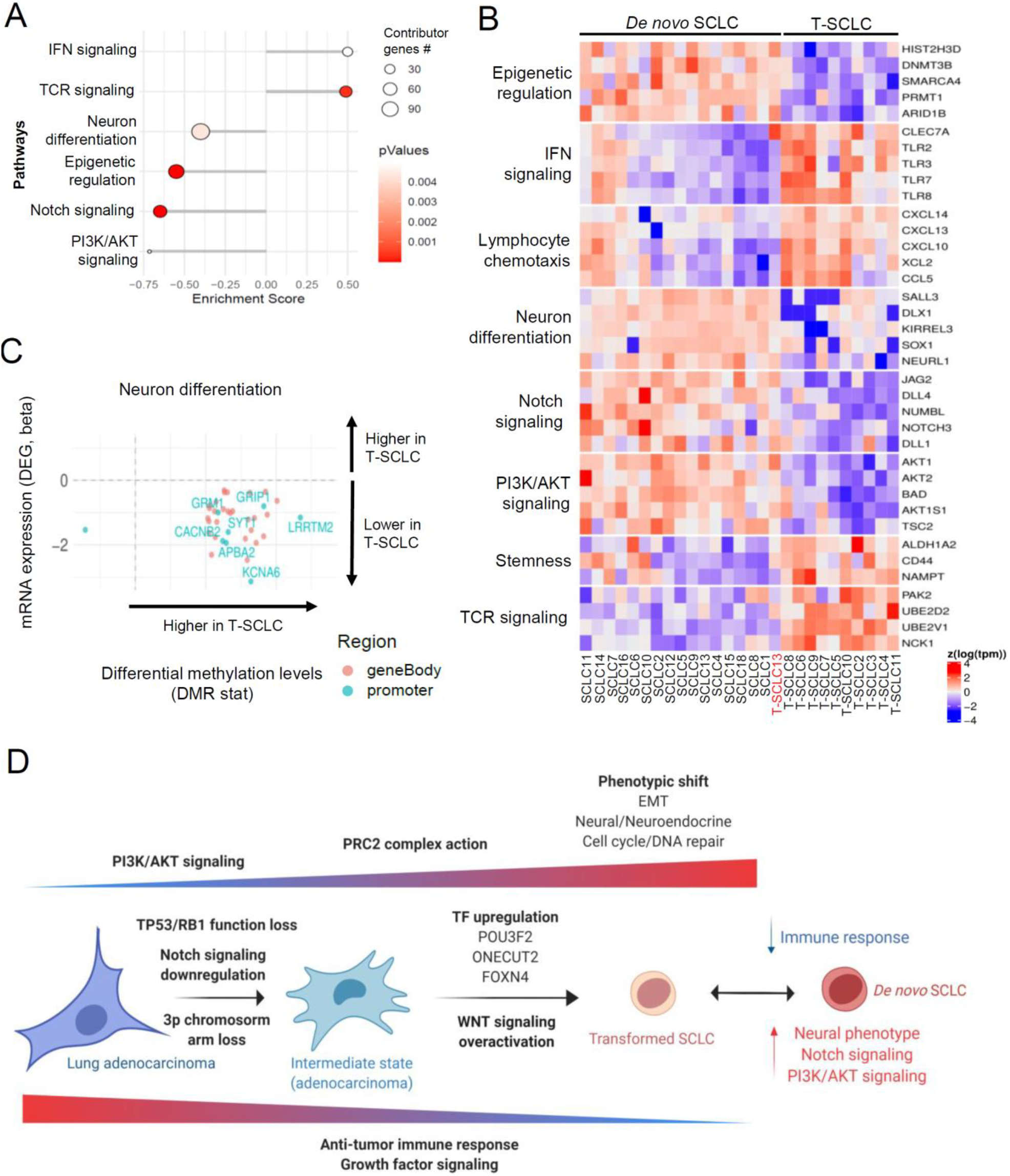
Integrative RNA and methylation analyses of T-SCLC versus *de novo* SCLC. Related to Supplementary Figure S7. (A) Pathway enrichment analyses on the DEGs of T-SCLC versus *de novo* SCLC comparison. (B) Heatmap highlighting DEGs of interest, grouped by recurrent pathways, of T-SCLC versus *de novo* SCLC comparison. (C) Scatter plots showing DEGs exhibiting differential methylation levels in T-SCLC versus *de novo* SCLC comparison, grouped by pathways of interest. Significantly differentially expressed (q value < 0.05 and beta >= log2(1.5)) and methylated (FDR < 0.5 and differential methylation level greater than 0.1) sites are highlighted. Those genes where increased gene body or promoter methylation is correlated to expression positively and negatively, respectively, are labeled. (D) Schematic of molecular and phenotype changes on the different steps of SCLC transformation. Our data suggest that transformation from LUAD to SCLC may be a progressive process involving multiple signaling pathways and phenotypic changes. This process may be initiated by the loss of *TP53* and *RB1*, decreased dependence on RTK signaling and Notch signaling downregulation, and involve progressive activation of AKT and WNT signaling pathways, epigenomic regulation by the PRC2 complex and a number of additional epigenetic enzymes, acquisition of a neuronal and EMT phenotype, and downregulation of genes involved in multiple immune response pathways. Samples IDs in black and red indicate that they come from a combined histology specimen or a pre-/post-transformation specimen, respectively.

## DISCUSSION

Cancer cell promiscuity in lineage commitment is a reflection of the exceptional heterogeneity of tumors, and an important source of treatment failure. The advent of potent and specific targeted inhibitors for mutational drivers in LUAD, like the use of highly effective anti-androgenic agents in prostate cancer, has prompted increasing recognition of lineage plasticity as a primary barrier to successful management of cancer. While frequently considered in the context of acquired therapeutic resistance, lineage plasticity in cancer is also evident independent of drug selection. In this study, we took advantage of the long-standing recognition of mixed histology lung cancers to gain insight into the molecular phenotypic landscapes underlying histologic transformation between LUAD and SCLC lineages. Whole exome sequencing confirmed that the histologically distinct components of mixed tumors were clonally related, reflecting distinct lineage pathways derived from a shared tumorigenic founder. By focusing primarily on a cohort of biphenotypic tumors in which the distinct lineages are in temporal and spatial proximity, we have the opportunity to identify consistent molecular changes that characterize this transformation. In this study, we provide the first comprehensive multi-omic characterization of NE transformation in lung cancer, including genomic, transcriptomic, epigenomic and proteomic analyses of matched samples.

One conclusion that may be taken from our data concerns the degree to which activation of lineage plasticity can result in distinct cell fates. *De novo* SCLC has been classified into four distinct subtypes based on differential expression of master transcriptional regulators^3,15,31^. Examining a cohort of mixed histology LUAD/SCLC tumors, we find that the T-SCLC derivatives do not consistently fall into one of these subtypes – rather we find all four subtypes clearly represented among just 11 cases. This observation underscores the degree to which plasticity in lung cancer can activate diverse transcriptional programs. Particularly surprising to us was the identification of mixed histology tumors in which the T-SCLC component expressed *POU2F3*, defining the subtype SCLC-P. LUAD is believed to derive from type II pneumocytes^32^. Based on its expression profile, SCLC-P had been proposed to arise from transformation of tuft cells, a rare pulmonary cell that is the exclusive source of *POU2F3* expression in lung^21^. The identification of two independent cases of clonally linked T-LUAD and *POU2F3*-expressing T-SCLC calls into question the cell of origin of SCLC-P and highlights the capacity of lineage plasticity to allow cancer cells to transdifferentiate between clearly distinct biological lineages.

Several features of the analysis of mixed histology T-LUAD/T-SCLC tumors reflect prior observations made regarding NE transformation of LUAD, reinforcing the relevance of this approach. Consistent with previous publications^9,13^, we observe inactivation of TP53 and RB1 by mutational or epigenetic mechanisms in essentially all T-SCLC, and in nearly all of the paired T-LUAD. This supports the role of concurrent *TP53/RB1* function loss as predictors of NE transformation^9^. While NE transformation of LUAD was originally observed in *EGFR*-mutant LUAD under selective pressure of EGFR TKI treatment, we confirm here similar histologic transformation regardless of EGFR mutation, including in the treatment naïve setting^3,8^. A novel finding here is the exceptionally high frequency of 3p chromosome arm loss in T-LUADs^33^. What genes resident on 3p singly or in combination could account for this observation is currently unclear, but 3p loss may represent a novel risk factor for NE transformation.

The paucity of recurrent mutations across samples in our cohort suggests that NE transformation in lung is not dependent on a common mutational driver, but rather may be primarily dependent on epigenetic shifts in gene expression programs. Transcriptional analysis of the T-LUAD and T-SCLC components of our mixed histology tumor set, relative to control (non-transformed) LUAD and *de novo* SCLC, suggests that T-LUADs and T-SCLCs occupy intermediate, transitional states – states that overlap both with their apparent non-transforming histology and with each other. Our data point to a number of signaling pathways that appear to shift in consistent patterns from T-LUAD to T-SCLC. These shifts include higher expression of genes in cell cycle and DNA repair, consistent with the highly proliferative capacity of SCLC tumors^34^. Higher expression of neuroendocrine and mesenchymal features in T-SCLC agrees with previous reports suggesting that NE transformation may occur through an intermediate EMT stem-like state^35,36^. Our data correlates this with putative methylation-induced repression of cell adhesion molecules, and induced expression of mesenchymal effectors such as *CDH2* (N-cadherin) and *NCAM1* associated with demethylation of binding motifs of key mediators of EMT, such as *SNAI1* and *TWIST1* in T-SCLC.

Our data implicates multiple pathways known to regulate stem and progenitor cell biology in lineage plasticity and NE transformation, notably including upregulation of PRC2 complex activity, induction of WNT signaling and suppression of the Notch pathway. The induction of PRC2 activity is in keeping with its apparent role in NE transformation in prostate cancer^4,37^. WNT signaling too has been previously implicated in lineage plasticity^38^ and in the maintenance of a NE phenotype in the prostate^39,40^. Given the previously defined role of Notch signaling in suppression of NE tumor growth, we believe that sustained inhibition of Notch signaling may be a prerequisite for NE transformation in lung.

We also find consistent evidence of PI3K/AKT pathway activation in T-SCLC. Emerging data supports a role for PI3K/AKT signaling in lineage plasticity and neuroendocrine transformation^3,41^. AKT also has been identified as a driver of NE phenotypic shift in non-tumoral prostate and lung cells^42^.

Finally, we note that SCLCs are notoriously immune “cold” tumors relative to NSCLCs^14,15,43^. Consistent with this, we see a progressive suppression in anti-tumor immune response pathways including cytokine signaling, T-cell immunity, and neutrophil degranulation from control LUAD to T-LUAD, from T-LUAD to T-SCLC, and from T-SCLC to *de novo* SCLC.

NE transformation in lung cancer induces a highly lethal and recalcitrant tumor profile that currently lacks effective treatments. We need better understanding of molecular drivers of NE transformation in lung cancer to identify therapeutic targets to treat or prevent transformation. Through detailed analysis of mixed histology pairs, we provide the first comprehensive molecular characterization of NE transformation in lung cancer, describing the signaling pathways and phenotypes altered during histologic transformation mediated by lineage plasticity. Notably, many of the pathways identified in this study, including PI3K/AKT, WNT, and EZH2, are druggable and are being actively targeted in ongoing clinical trials. The development of representative preclinical models of NE transformation in human lung cancer remains a primary unmet need: development and interrogation of these pathways in such models could further inform prevention or intervention strategies for disruption of lineage plasticity in lung cancer patients.

## METHODS

### Clinical samples

We identified 11 formalin-fixed paraffin-embedded (FFPE) tumors with combined LUAD and SCLC histology, from which independent isolation of both histological components was possible (N=11, **Supplementary Tables S1-2, Supplementary Figure S1**). As the components of these mixed histology tumors are not temporally ordered, we refer to the component parts of these mixed histology tumors as “T-LUAD” and “T-SCLC” with the T referring to histologic transformation. We identified an additional 5 pre-transformation LUAD and 3 post-transformation SCLC cases for which tissue material was available (**Supplementary Tables S1, S3**). As controls we included a group of never-transformed LUADs (N=15) and a set of *de novo* SCLC samples (N=18) (**Supplementary Tables S1, S4**). All study subjects had provided signed informed consent for biospecimen analyses under an Institutional Review Board-approved protocol.

### Tissue isolation

For microdissection, hematoxylin and eosin (H&E)-stained FFPE tumor slides of tumors with combined LUAD/SCLC were independently evaluated by two pathologists. Where possible, multiple FFPE blocks of each tumor were reviewed, with the aim of selecting areas containing exclusively the LUAD or the SCLC component. Where individual slides with pure components were not available, slides containing both histologic components with complete physical separation were selected. Between 10 and 20 unstained sections (USS) at 10μm prepared on uncharged slides from corresponding FFPE blocks were used for microdissection of each case. Every 10 sections, an additional section was stained with H&E for confirmation of histology. The areas corresponding to each histological component on the initial H&E were dissected using a clean blade and the tissue collected in 0.5ml nuclease free tubes for nucleic acid extraction. Alternatively, 1.0-1.5mm core punches were made from LUAD and SCLC areas on the FFPE blocks and placed in 0.5ml nuclease free tubes for nucleic acid extraction, exclusively in cases where each histologic component was located in a different block, and where no histologic cross-contamination was confirmed by pathological review.

### DNA Extraction

FFPE tissue was deparaffinized using heat treatment (90°C for 10’ in 480μL PBS and 20μL 10% Tween 20), centrifugation (10,000x*g* for 15’), and ice chill. Paraffin and supernatant were removed, and the pellet was washed with 1mL 100% EtOH followed by an incubation overnight in 400μl 1M NaSCN for rehydration and impurity removal. Tissues were subsequently digested with 40μl Proteinase K (600 mAU/ml) in 360μl Buffer ATL at 55°C. DNA isolation proceeded with the DNeasy Blood & Tissue Kit (QIAGEN catalog # 69504) according to the manufacturer’s protocol modified by replacing AW2 buffer with 80% ethanol. DNA was eluted in 0.5X Buffer AE.

### RNA/DNA dual extraction from FFPE tissue

FFPE sections were deparaffinized in mineral oil. Briefly, 800μL mineral oil (Fisher Scientific, #AC415080010) and 180μL Buffer PKD were mixed with the sections, Proteinase K was added for tissue digestion, and the sample was incubated at 56°C for 15 minutes. Phase separation was encouraged with centrifugation, and the aqueous phase was chilled 3 minutes to precipitate RNA. After centrifugation for 15 minutes at 20,000*g*, RNA-containing supernatant was removed for extraction, while DNA remained in the pellet. Nucleic acids were subsequently extracted using the AllPrep DNA/RNA Mini Kit (QIAGEN, #80204) according to the manufacturer’s instructions. RNA was eluted in nuclease-free water and DNA in 0.5X Buffer ATE.

### RNA/DNA dual extraction from frozen tissue

Frozen tissues were weighed and homogenized in RLT and nucleic acids were extracted using the AllPrep DNA/RNA Mini Kit (QIAGEN, #80204) according to the manufacturer’s instructions. RNA was eluted in nuclease-free water and DNA in 0.5X Buffer EB.

### Whole exome sequencing from DNA

After PicoGreen quantification and quality control by Agilent BioAnalyzer, 100-500 ng of DNA were used to prepare libraries using the KAPA Hyper Prep Kit (Kapa Biosystems KK8504) with 8 cycles of PCR. After sample barcoding, 100 ng of library were captured by hybridization using the xGen Exome Research Panel v1.0 (IDT) according to the manufacturer’s protocol. PCR amplification of the post-capture libraries was carried out for 12 cycles. Samples were run on a HiSeq 4000 in a 100bp/100bp paired end run, using the HiSeq 3000/4000 SBS Kit (Illumina). Normal and tumor samples were covered to an average of 66X and 76X, respectively.

### Whole exome sequencing from previous DNA libraries

After PicoGreen quantification and quality control by Agilent BioAnalyzer, 100 ng of library transferred from the DMP were captured by hybridization using the xGen Exome Research Panel v1.0 (IDT) according to the manufacturer’s protocol. PCR amplification of the post-capture libraries was carried out for 8 cycles. Samples were run on a HiSeq 4000 in a 100bp/100bp paired end run, using the HiSeq 3000/4000 SBS Kit (Illumina). Normal and tumor samples were covered to an average of 114X and 202X, respectively.

### Whole Exome Analysis

We used a comprehensive in-house WES pipeline TEMPO - Time efficient mutational profiling in oncology (https://github.com/mskcc/tempo) that performs alignment using BWA-mem algorithm followed by mutation calling using Strekla2 and Mutect2 variant callers. The combined, annotated and filtered variant calls were used for downstream analysis. Details of the variant call processing are described at https://ccstempo.netlify.com/variant-annotation-and-filtering.html#somatic-snvs-and-indels and are previously described as well^44^. Copy-number analysis was performed with FACETS (https://github.com/mskcc/facets), processed using facets-suite (https://github.com/mskcc/facets-suite), and manual reviewed and refitted using facets-preview (https://github.com/taylor-lab/facets-preview). To delineate mutational processes driving the acquisition of somatic alterations, mutational signatures were decomposed for all tumor samples that had a minimum of 5 single-nucleotide somatic mutations using the R package mutation-signatures (https://github.com/mskcc/mutation-signatures). Further, a given signature was considered to be ‘dominant’ if the proportion of mutations contributing to the signature was at least 20% of all mutations detected in the sample.

Purity, ploidy, tumor mutational burden (TBM), genome doubling, and cancer cell fractions for all mutations in all specimens were inferred from sequencing data. We estimated neoantigen load by taking the number of variant estimated to having strong class I MHC binding affinity by NetMHC 4.0^45^ and normalizing it by the TMB. We summarized the top occurring somatic variants located on cancer genes in an oncoprint using the R package ComplexHeatmaps version 2.0.0 (https://github.com/jokergoo/ComplexHeatmap)^46^. Cancer genes were genes defined as “OncoKB Annotated” on the Cancer Gene List downloaded on June 2020 (https://www.oncokb.org/cancerGenes). All other plots for this analysis were created using ggplot version 3.3.2 (https://github.com/tidyverse/ggplot2).

### Comparison to TCGA

Somatic mutations and copy number alterations (CNAs) found in cancer genes in our T-LUAD samples were compared to those found in The Cancer Genome Atlas Lung Adenocarcinoma (TCGA-LUAD) cohort using a Fisher exact test. The mutations from TCGA-LUAD^47^ were extracted using the R package TCGA mutations (https://github.com/PoisonAlien/TCGAmutations) and tested against our cohort mutations with maftools v.2.0.16 (https://github.com/PoisonAlien/maftools)^48^. Separately, a Fisher exact test was used to identify significant CNAs by comparing the number of samples with amplifications and deletions on particular genes in TCGA-LUAD, extracted from CbioPortal^49,50^, to the number of samples with gene level CNAs in our cohort. For both mutations and CNAs, genes with p<0.05 were considered differentially altered. Lastly, the number of samples with 3p arm level loss in TCGA, extracted from CbioPortal, was compared the number of T-LUAD samples identified using FACETS with the same loss. Significance was identified using a Fisher exact test. The results were summarized in a volcano plot using the R packages, EnhancedVolcano version 1.7.4 (https://github.com/kevinblighe/EnhancedVolcano) and ggplot.

### Genetic Evolution

We estimated the clonal history for the combined histology cases with sufficient purity (>0.3) in both their T-LUAD and T-SCLC components. We first genotyped all somatic single nucleotide polymorphisms, located on cancers gene, that were called, in either T-LUAD or T-SCLC samples, in both tumor specimens per case. Genotyping was performed using GetBaseCountsMultiSample v.1.2.2 (https://github.com/mskcc/GetBaseCountsMultiSample). Using the new mutant allele fractions, cancer cell fraction and clonality for these mutations were inferred by the ccf-annotate-maf function from facets-suite, a process that has been previously described^44^. Mutations that were estimated to be clonal in both the T-LUAD and T-SCLC specimens were categorized as truncal mutations. Mutations that were clonal in only one specimen were classified to represent that branch of the clonal tree. The evolutionary trees were drawn manually with the length of each branch drawn proportionally to the number of clonal mutations.

The relative timing of mutations with respected to a global whole genome doubling (WGD) event was inferred as previously described^51^. In short, the most parsimonious explanation of an observed copy number state was used. Additionally, the observed copy number for all the segments for both the mutant and minor allele were calculated by FACETs and plotted using custom code to show common chromosomal copy number gains and losses.

### Methylation sequencing

After PicoGreen quantification (ThermoFisher, #P11496) and quality control by Agilent BioAnalyzer, 170-750 ng of genomic DNA were sheared using a LE220-plus Focused-ultrasonicator (Covaris, #500569). Samples were cleaned using Sample Purification Beads from the TruSeq Methyl Capture EPIC LT Library Prep Kit (Illumina, #FC-151-1002) according to the manufacturer’s instructions with modifications. Briefly, samples were incubated for 5 minutes after addition of SPB, 50 μL RSB were added for resuspension, and resuspended samples were incubated for 2 minutes. Sequencing libraries were prepared using the KAPA Hyper Prep Kit (Kapa Biosystems KK8504) without PCR amplification. Post-ligation cleanup proceeded according to Illumina’s instructions with 110 μL Sample Purification Mix. After purification, 3-4 samples were pooled equimolar and methylome regions were captured using EPIC oligos. Capture pools were bisulfite converted and amplified with 11-12 cycles of PCR. Pools were sequenced on a NovaSeq 6000 or HiSeq 4000 in a 150/150bp or 100bp/100bp paired end run, using the NovaSeq 6000 S4 Reagent Kit (300 Cycles) or HiSeq 3000/4000 SBS Kit (Illumina). The average number of read pairs per sample was 51 million.

### DNA methyl capture EPIC data processing

The Bismark pipeline^52^ was adopted to map bisulfite treated EPIC sequencing reads and determine cytosine methylation states. Trim Galore v0.6.4 was used to remove raw reads with low-quality (less than 20) and adapter sequences. The trimmed sequence reads were C(G) to T(A) converted and mapped to similarly converted reference human genome (hg19)^53^ using default Bowtie 2^54^ settings within Bismark. Duplicated reads were discarded. The remaining alignments were then used for cytosine methylation calling by Bismark methylation extractor.

### Differential methylation analysis

Differentially methylated CpGs (DMCs) were identified using DSS R package^55,56^ on the basis of dispersion shrinkage followed by Wald statistical test for beta-binomial distributions. Any CpGs with FDR < 0.05 and methylation percentage difference greater than 10% were considered significant DMCs. Differentially methylated regions (DMRs) were subsequently called based on the DMCs. The called DMRs were required to satisfy the minimum length of 50bps and minimum 3 CpGs in the region; two neighboring DMRs were merged if less than 50bps apart; and significant CpGs were those that occupy at least 50% of all CpGs population in the called DMRs as default in DSS package. Pairwise comparisons were conducted for pre-transformation LUAD vs control LUAD, post-transformation SCLC vs *de novo* SCLC, and post-transformation SCLC vs pre-transformation LUAD. The DMRs were mapped to gene regions at promoters and gene bodies, and differential methylation levels were subsequently associated with differential gene expression values in selected pathways. In addition to pairwise comparisons, principal component analysis (PCA) and partial least square discriminant analysis (PLSDA) were also performed to classify samples into groups and identify influential CpGs using mixOmics R package^56^.

### Motif enrichment analysis

Differential methylation may influence transcription factor (TF) binding. To identify overrepresented known TF motifs due to differential methylation for the post-transformation SCLC compared with pre-transformation LUAD, “findMotifsGenome.pl” from HOMER^57^ was applied to DMCs (+/−50bps) overlapping with gene promoter regions. DMCs regions with hyper- and hypo-methylation in SCLC were explored separately to show the effects from different methylation status. The significantly enriched TFs were defined as those with p value ≤ 0.05.

### RNA sequencing

Approximately 500ng of FFPE RNA or 100ng of fresh frozen RNA per sample were used for RNA library construction using the KAPA RNA Hyper library prep kit (Roche, Switzerland) per the manufacturer’s instructions with minor modifications. Customized adapters with unique molecular indexes (UMI) (Integrated DNA Technologies, US) and Sample-specific dual-indexes primers (Integrated DNA Technologies, US) were added to each library. The quantity of libraries was measured with Qubit (Thermo Fisher Scientific, US) and quality measured by TapStation Genomic DNA Assay (Agilent Technologies, US). Equal amounts of each RNA library (around 500ng) were pooled for hybridization capture with IDT Whole Exome Panel V1 (Integrated DNA Technologies, US) using a customized capture protocol modified from NimbleGen SeqCap Target Enrichment system (Roche, Switzerland). The captured DNA libraries were then sequenced on an Illumina HiSeq4000 with paired end reads (2Å~100bp), at 50millions reads/sample.

### RNASeq Analysis

In-line UMI sequences were trimmed from the sequencing reads with Marianas (https://github.com/mskcc/Marianas) and aligned to human GRCh37 genome using STAR 2.7.0 (https://github.com/alexdobin/STAR)^58^ with Ensembl v75 gene annotation. Hybrid selection specific metrics and Alignment metrics were calculated for the BAM files using CalculateHsMetrics and CollectRnaSeqMetrics, respectively, from Picard Toolkit (https://github.com/broadinstitute/picard) to determine the quality of the capture.

We quantified RNA-seq reads with Kallisto v.0.45.0^59^ to obtain transcript counts and abundances. Kallisto was run with 100 bootstrap samples, sequence based bias correction, and in strand specific mode, which processed only the fragments where the first read in a pair is pseudoaligned to the reverse strand of a transcript. Differential gene expression analysis, principle component analysis, and transcript per million (TPM) normalization by size factors, were done from Kallisto output files using Sleuth v0.30.0 run in gene mode^60^. Differentially expressed genes were identified using the Wald test. Genes were marked significant if the False Discovery Rates, *q*, calculated using the Benjamini-Hochberg menthod, was less than 0.05, and *beta*(Sleuth-based estimation of log2 fold change)>1.25, which approximately correlated to a log2 fold change of 2 in our data. The log of the normalized TPM values for selected significant genes, were rescaled using a z-score transformation, and plotted in a heatmap using the ComplexHeatmap Library in R.

### Pathway enrichment

Gene set enrichment analysis (GSEA)^61^ was performed on full sets of gene expression data across the previously mentioned three comparisons. Genes were ranked on p value scores computed as −log10(p value)*(sign of beta). Gene set annotations were taken from Molecular Signatures Database (MSigDB v7.0.1)^61,62^. Gene sets tagged by KEGG^63,64^ and REACTOME^65^ pathways were retained for further analysis. The significance level of enrichment was evaluated using permutation test and the p value was adjusted by Benjamini-Hochberg procedure. Any enriched gene sets with adjusted p value ≤ 0.05 were regarded as significant. This analysis was conducted using ClusterProfiler R package^66^. The enriched gene sets that are influenced by DMCs were selected and pathway annotations concatenated manually to remove redundancy and achieve high level generality. When the pathway terms were merged, median enrichment score was taken as the new group enrichment score, p values were aggregated using Fisher’s method from the Aggregation R package^67^, and core enrichment of genes were collapsed.

### Phospho-kinase array

Protein samples were quantified with the Bradford method (#5000205, Bio-Rad) and 200 ug aliquots were used in the phospho-kinase array (#ARYC003C, R&D-Biotechne), which was performed using the manufacturer’s instructions. Quantification of spots was performed using the Image Studio software (Version 3.1, Li-Cor). Technical replicates (2 per array) per sample were averaged. Two-tailed Student’s T-test was performed on these values, comparing the T-LUAD and T-LUSC groups.

### Cell line transductions

PC9 cell line was purchased from Millipore Sigma (#90071810-VL) and HCC827 cell line was purchased from ATCC (#CRL-2868). Both cell lines were regularly tested for Mycoplasma and maintained in RPMI 1640 10% FBS. Lentiviruses were produced as previously described^68^ with FOXN4 (#EX-I2262-Lv151, GeneCopoeia), POU3F2 (#EX-A3238-Lv151, GeneCopoeia) and ONECUT2 (#EX-Z4476-Lv151, GeneCopoeia) overexpression lentiviral plasmids, with a EGFP overexpression plasmid as control plasmid (#EX-EGFP-Lv151, Genecopoeia). Cell lines were transduced at high MOI as previously described^68^ with overnight virus incubation.

### Immunoblotting

Protein extraction and western blot were performed as previously described^69^. Antibodies for FOXN4 (#PA539174, ThermoFisher), ONECUT2 (#ab28466, Abcam), POU3F2 (#12137, Cell Signaling Technology), EGFR (#4267, Cell Signaling Technology) and actin (#3700, Cell Signaling Technology) were used.

### RT-qPCR

RNA extraction, reverse transcription and quantitative PCR were performed as previously described^70^. *FOXN4* expression was normalized to that of *GAPDH*. Fluorescent probes against *FOXN4* (#4351372, Applied Biosystems) and GAPDH (#4331182, Applied Biosystems) were used.

## ACKNOWLEDGEMENTS

Supported by NCI R01 CA197936 and U24 CA213274 (CMR), the SU2C/VAI Epigenetics Dream Team (CMR), the Druckenmiller Center for Lung Cancer Research (CMR, TS, AQV), Parker Institute for Cancer Immunotherapy grant (TS); International Association for the Study of Lung Cancer grant (TS), NIH K08 CA-248723 (AC). We acknowledge the use of the Integrated Genomics Operation Core, funded by the NCI Cancer Center Support Grant (CCSG, P30 CA08748), Cycle for Survival, and the Marie-Josée and Henry R. Kravis Center for Molecular Oncology. We also acknowledge Maria Corazon Mariana and Emily Lin from the PPBC Biobank for their invaluable help. The PPBC Biobank and Pathology Core Facility are supported by the NCI Cancer Center Support Grant P30-CA008748.

