## Supplemental Figures and Figure Legends for "Multi-omic analysis of lung tumors defines pathways activated in neuroendocrine transformation"

### SUPPLEMENTARY FIGURES AND FIGURE LEGENDS

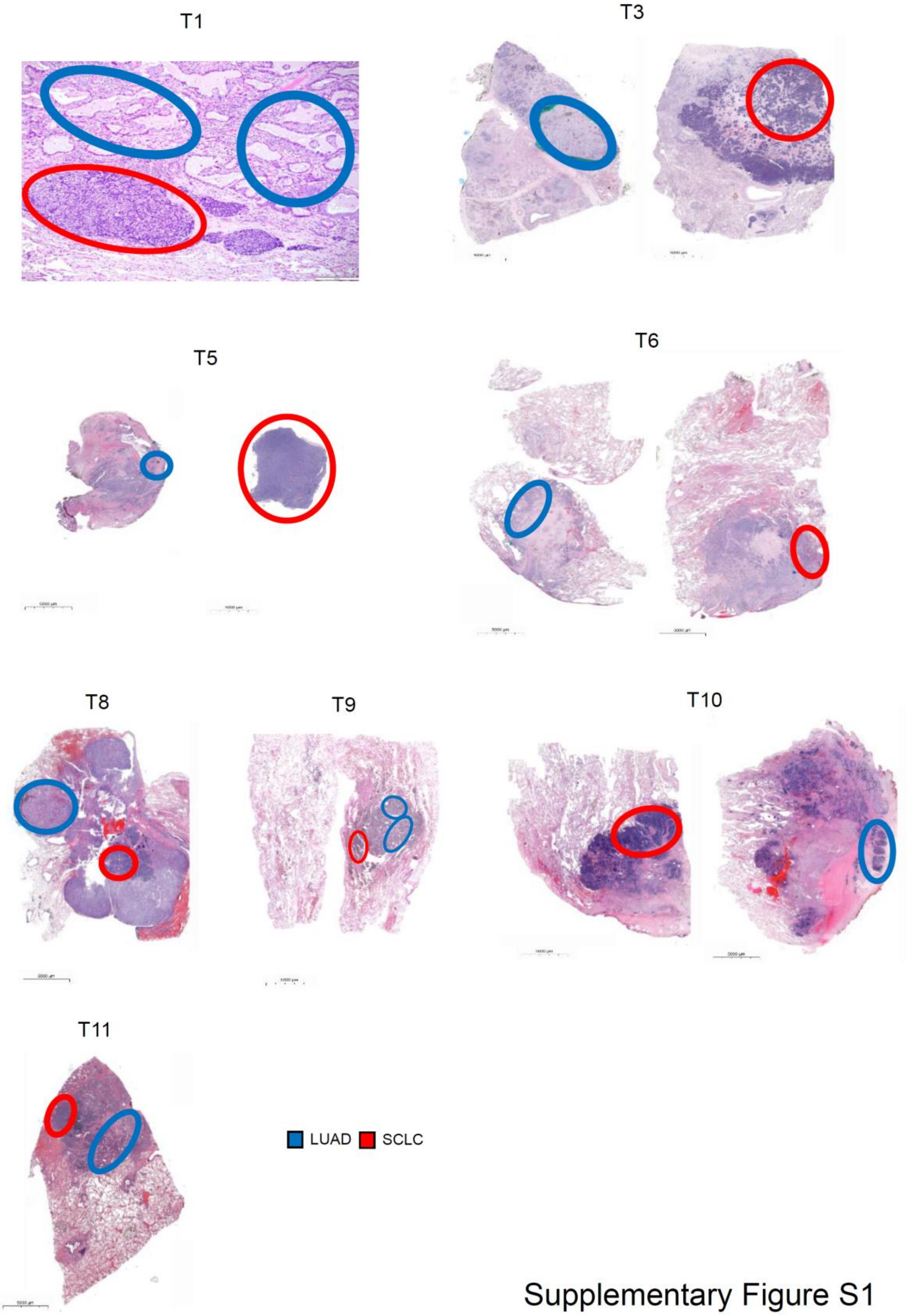


**Supplementary Figure S1.** H&E staining of the combined LUAD/SCLC cases in our cohort with histologic components labeled.

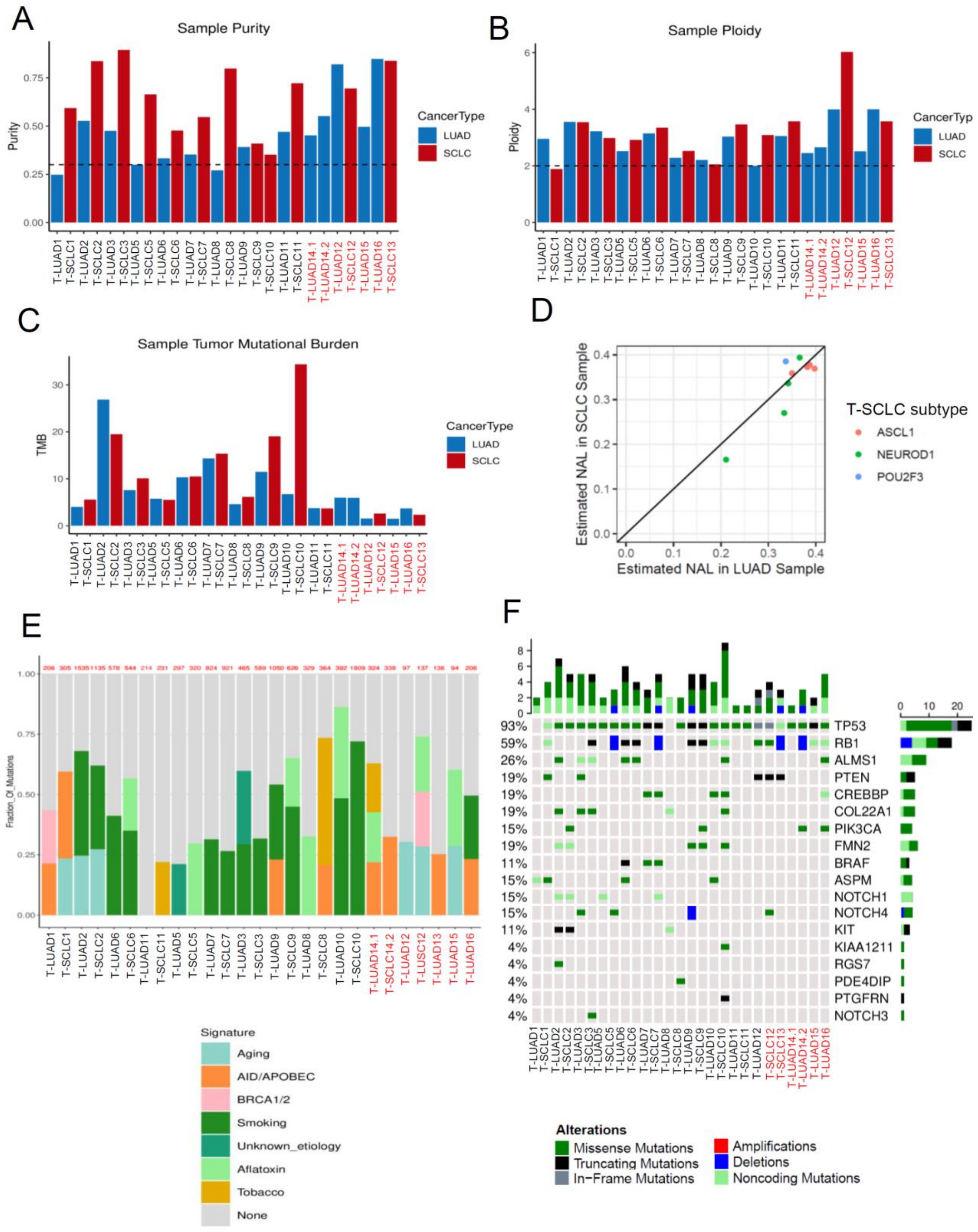


5

6 **Supplementary Figure S2.** Bar plots showing tumor purity (A), ploidy (B) and mutational

burden (C) of samples in the cohort analyzed by WES. (D) Neoantigen load in matched T-LUAD and T-SCLC samples labeled by subtype of the SCLC component. (E) Enrichment in mutational signatures on the samples in our cohort analyzed by WES. (F) Oncoprint of mutations frequently found in T-SCLC[71] in our samples analyzed by WES. Samples IDs in black and red indicate that they come from a combined histology specimen or a pre-/post-transformation specimen, respectively.

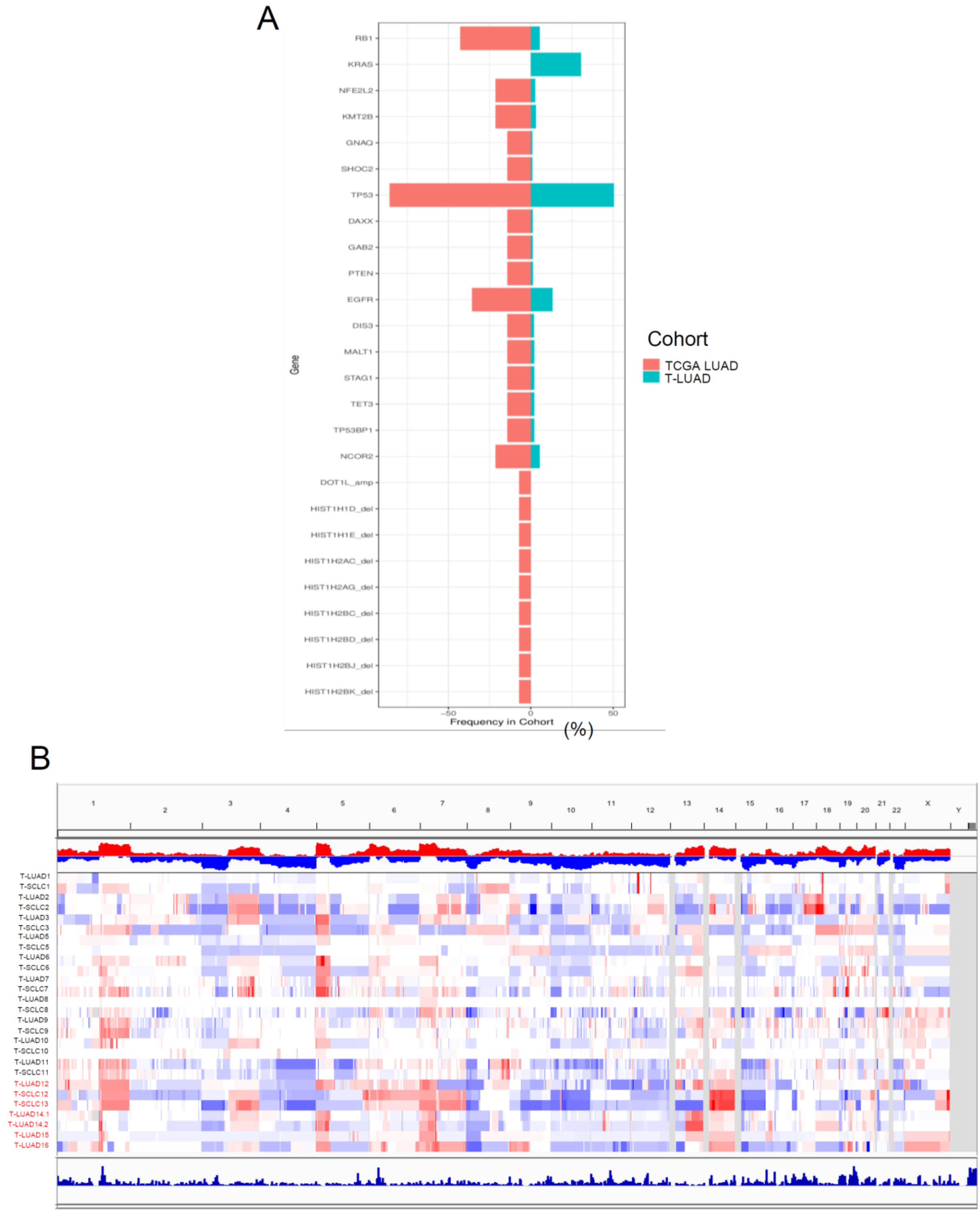


Supplementary Figure S3

**Supplementary Figure S3**. (A) Prevalence (%) of mutations and CNAs enriched in our cohort of T-LUADs versus TCGA LUAD cohort (p-value<0.05). (B) Chromosomal amplifications/deletions in samples in the transformation cohort analyzed by WES. Samples IDs in black and red indicate that they come from a combined histology specimen or a pre-/post-transformation specimen, respectively.

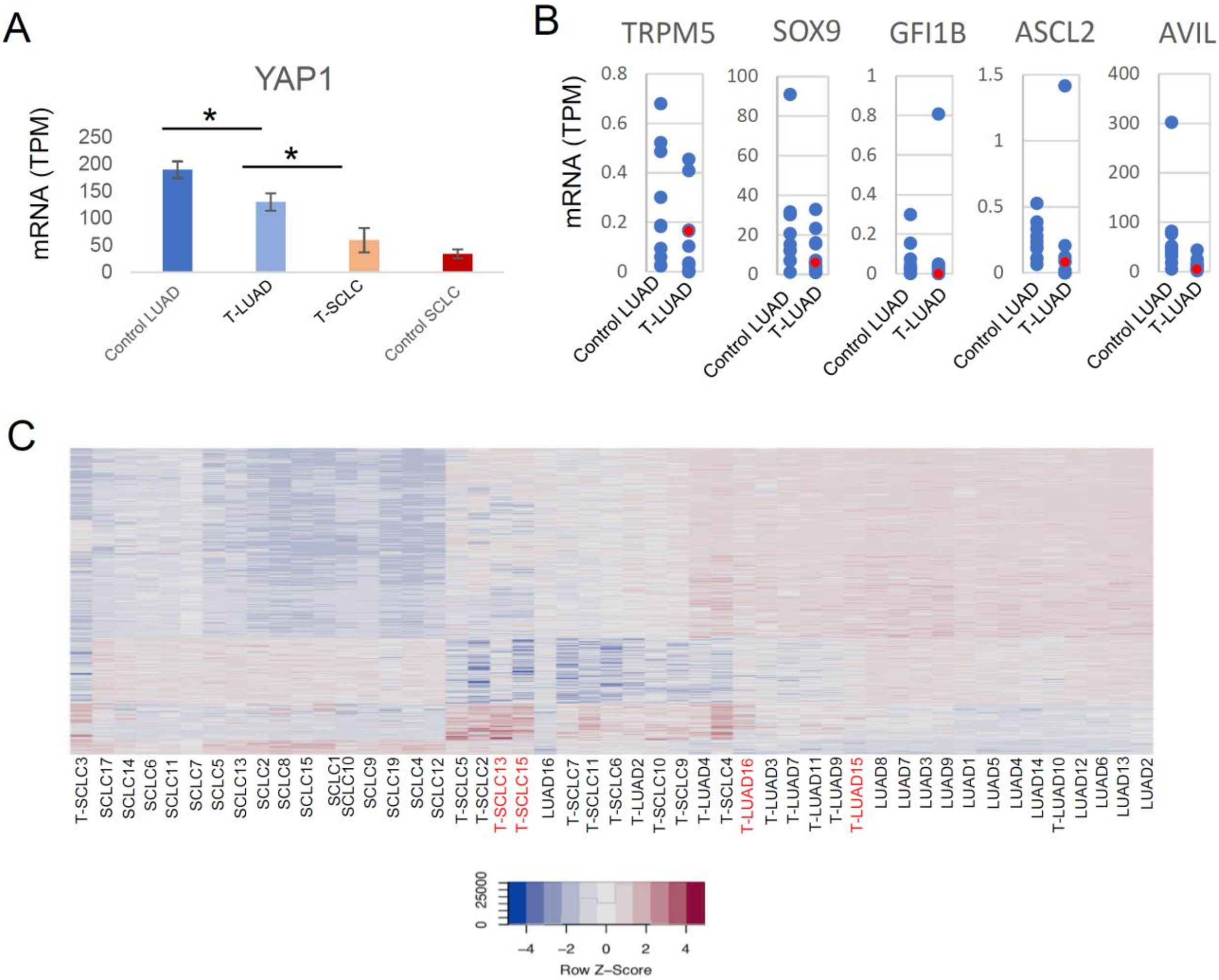


**Supplementary Figure S4.** (A) YAP1 mRNA expression in control LUAD, T-LUAD, T-SCLC, and *de novo* SCLC samples. Two tailed Student's t-test was used to assess statistical significance of the differential expression between groups (B) mRNA expression of tuft cell markers[21] in our control and T-LUADs. The expression values for the LUAD component of T3 are highlighted in red. (C) Heatmap of methylation levels on PLSDA components 1 and 2. Samples IDs in black and red indicate that they come from a combined histology specimen or a pre-/post-transformation specimen, respectively. p-values legend: * $p<0.05$.

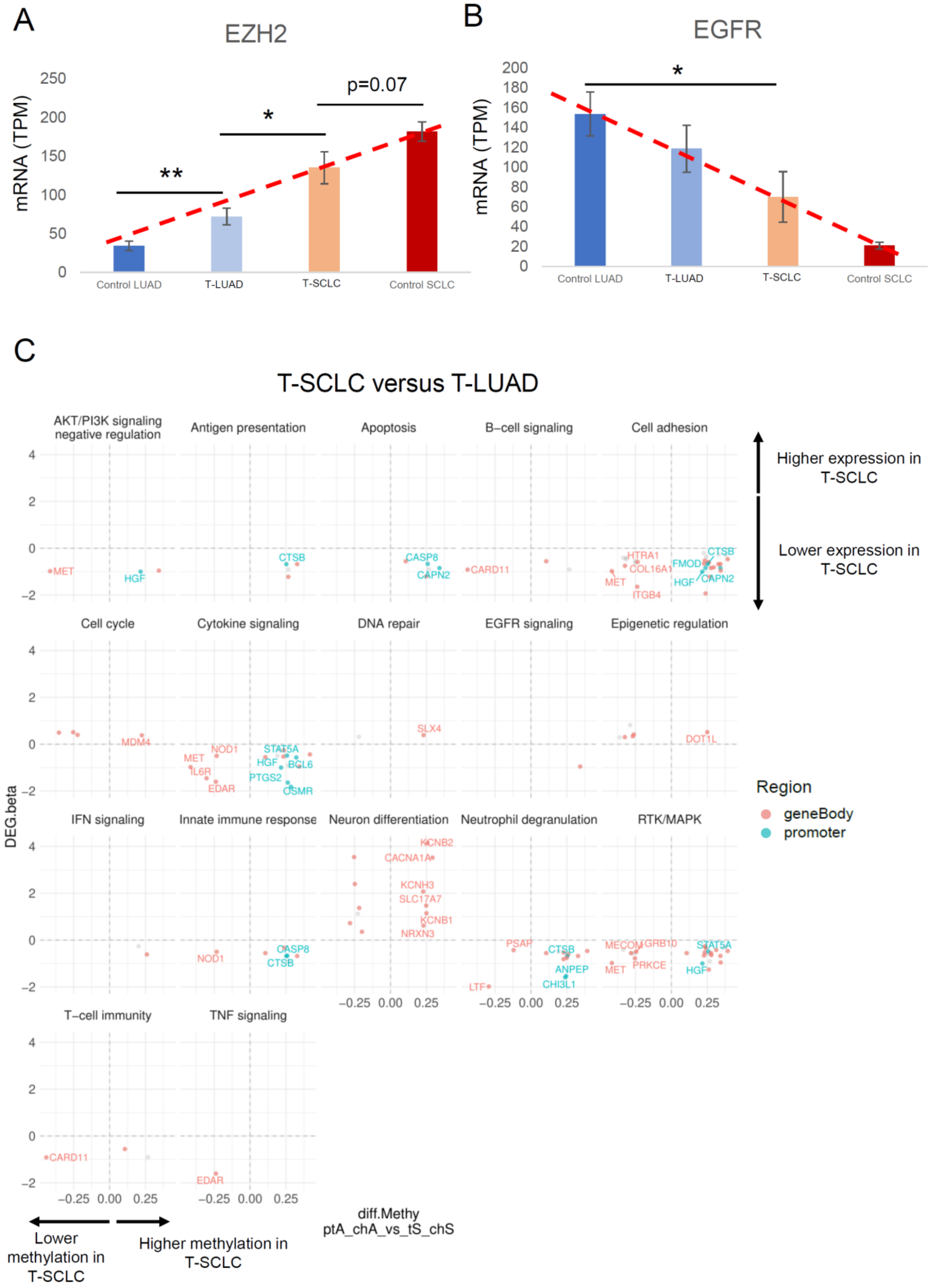


83 ## Supplementary Figure S5

84 **Supplementary Figure S5.** *EZH2* (A) and *EGFR* (B) mRNA expression in control LUAD,

T-LUAD, T-SCLC, and *de novo* SCLC samples. Two tailed Student's t-test was used to assess statistical significance of the differential expression between groups (C) Scatter plots showing DEGs exhibiting differential methylation levels in T-LUAD vs. T-SCLC, grouped by pathways found enriched or depleted in our pathway enrichment analyses on RNA data. For DEGs, beta value (Sleuth-based estimation of log2 fold change) is shown. Significantly differentially expressed (q value < 0.05 and beta >= log2(1.5)) and methylated (FDR < 0.5 and differential methylation level greater than 0.1) sites are highlighted. Those genes where increased gene body or promoter methylation is correlated to expression positively and negatively, respectively, are labeled. p-values legend: * p<0.05, **p<0.01.

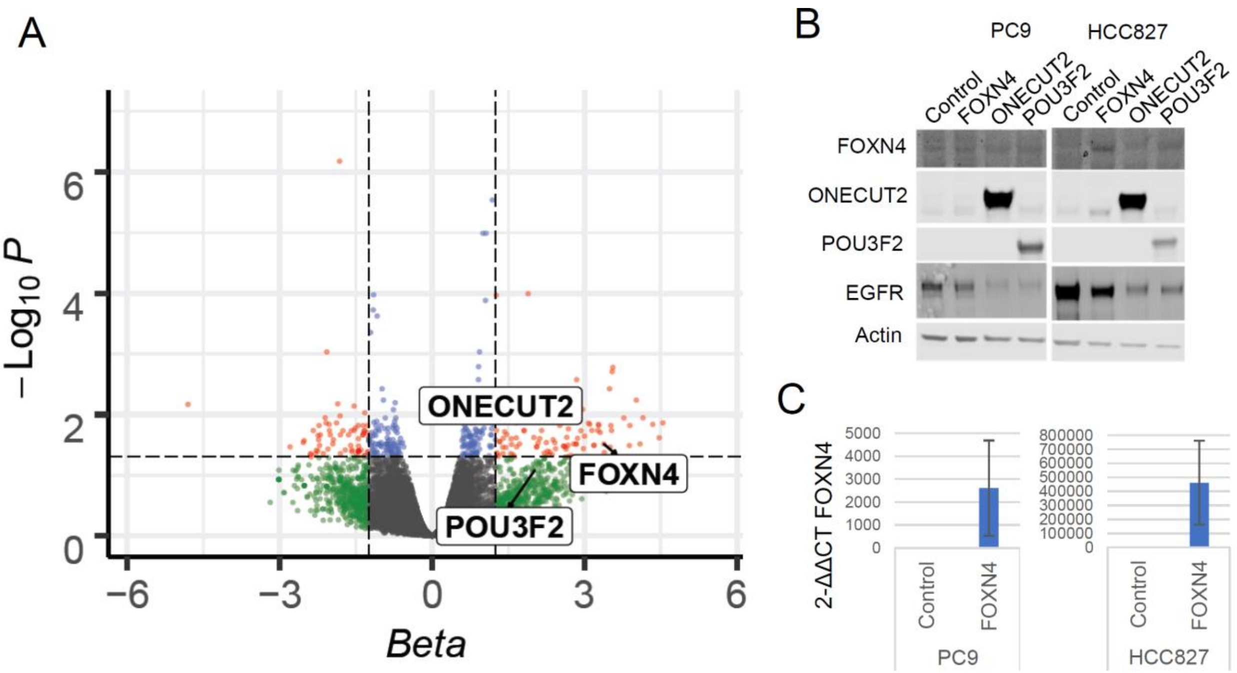


**Supplementary Figure S6.** (A) Volcano plot showing overexpression of transcription factors of interest at the RNA level in T-SCLC versus T-LUAD. For DEGs, beta value (Sleuth-based estimation of log2 fold change) is shown. (B) Western blots exhibiting downregulation of EGFR after overexpression of FOXN4, ONECUT2 or POU3F2 in two *EGFR*-mutant lung LUAD cell lines. (C) Confirmation of FOXN4 overexpression at the RNA level by qPCR.

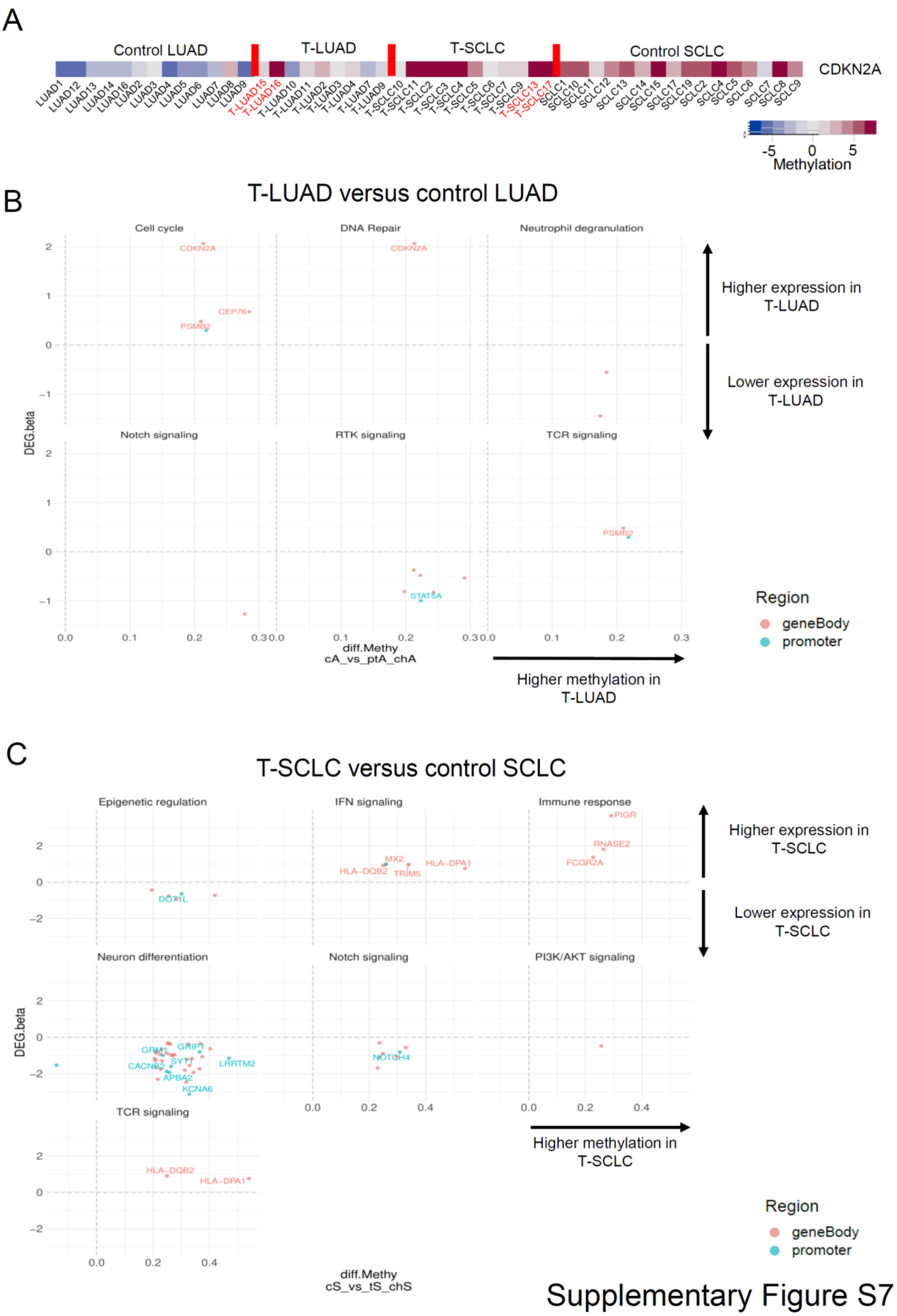


**Supplementary Figure S7**. (A) *CDKN2A* body methylation levels in the samples in our cohort analyzed by EPIC. Scatter plots showing DEGs exhibiting differential methylation levels in our T-LUAD versus control LUAD (B) or T-SCLC versus *de novo* SCLC (C) comparisons, grouped by pathways enriched or depleted in pathway enrichment analyses on RNA. For DEGs, beta value (Sleuth-based estimation of log2 fold change) is shown. Significantly differentially expressed (q value < 0.05 and beta >= log2(1.5)) and methylated (FDR < 0.5 and differential methylation level greater than 0.1) sites are highlighted. Those genes where increased gene body or promoter methylation is correlated to expression positively and negatively, respectively, are labeled. Samples IDs in black and red indicate that they come from a combined histology specimen or a pre-/post-transformation specimen, respectively.
